# Decoding Brain Structure-Function Dynamics in Health and in Psychosis: A Tale of Two Models

**DOI:** 10.1101/2024.10.03.616264

**Authors:** Qing Cai, Hannah Thomas, Vanessa Hyde, Pedro Luque Laguna, Carolyn B. McNabb, Krish D. Singh, Derek K. Jones, Eirini Messaritaki

## Abstract

Understanding the intricate relationship between brain structure and function is a cornerstone challenge in neuroscience, critical for deciphering the mechanisms that underlie healthy and pathological brain function. In this work, we present a comprehensive framework for mapping structural connectivity measured via diffusion-MRI to resting-state functional connectivity measured via magnetoencephalography, utilizing a deep-learning model based on a Graph Multi-Head Attention AutoEncoder. We compare the results to those from an analytical model that utilizes shortest-path-length and search-information communication mechanisms. The deep-learning model outperformed the analytical model in predicting functional connectivity in healthy participants at the individual level, achieving mean correlation coefficients higher than 0.8 in the alpha and beta frequency bands. Our results imply that human brain structural connectivity and electrophysiological functional connectivity are tightly coupled. The two models suggested distinct structure-function coupling in people with psychosis compared to healthy participants (*p* < 2 × 10^−4^ for the deep-learning model, *p* < 3 × 10^−3^ in the delta band for the analytical model). Importantly, the alterations in the structure-function relationship were much more pronounced than any structure-specific or function-specific alterations observed in the psychosis participants. The findings demonstrate that analytical algorithms effectively model communication between brain areas in psychosis patients within the delta and theta bands, whereas more sophisticated models are necessary to capture the dynamics in the alpha and beta band.

## 1 Introduction

Quantifying the mapping between structural connectivity (SC) and functional connectivity (FC) in the human brain is a major goal of neuroscience research^1^. Studying the SC-FC relationship in healthy and patient populations can reveal the origins and evolution of neurological disorders. It can enhance our understanding of the structural underpinnings of functional disorders, shedding light on whether such conditions primarily arise from structural damage of functional impairments. Current neuroimaging techniques, such as magnetic resonance imaging (MRI), functional MRI (fMRI), electroencephalography (EEG) and magnetoencephalography (MEG), allow for the non-invasive measurement of SC and FC.

Various methodologies have been used to relate SC to fMRI-measured FC^2–5^, including forward-generating models that link the brain’s structure to FC^6–9^, and analytical models that assume specific communication mechanisms between brain areas (such as shortest path, search information, navigation, etc.)^10^. These studies confirm that interactions among brain areas are influenced by the broad network of white matter connections in addition to shortest paths. SC has also been linked to elecrophysiologically-measured FC^11–16^, including via the use of analytical models^17^. This is of particular interest, given that electrophysiological FC reflects brain connectivity at the milli-second time scale, i.e., the actual time scale of functional interactions in the human brain.

Even though analytical models capture a significant fraction of the SC-FC relationship and elucidate the mechanisms via which communication happens between brain areas, they cannot be guaranteed to capture all aspects of their complex association. Machine-learning approaches, on the other hand, are well-suited to this kind of complexity, as recently shown for fMRI-measured FC^18^. To effectively link the complex nature of structural and functional connectivity patterns, we require a model that can naturally express these connectivity relationships. Graph Neural Networks (GNNs) are specifically designed for this purpose. GNNs are neural networks tailored for graph-structured data, where nodes represent entities and connections (edges) denote the relationships between them^19^. GNNs excel at handling these graph structures by iteratively aggregating information from neighboring nodes, thereby creating enriched representations of each node. This process captures the complex interactions within the graph and enhances the model’s understanding of the underlying relationships.

In this work we investigate for the first time how well a GNN model can predict electrophysiological resting-state functional connectivity from structural connectivity. We leverage node attributes and local topological information derived from SC matrices to generate high-dimensional representations that capture the properties of the underlying connectome. To achieve this, we employ a Graph Multi-Head Attention Autoencoder (GMHA-AE) based on multi-head attention mechanisms. Autoencoders, first introduced in 1986^20^, are neural networks that consist of an encoder and a decoder. The encoder compresses the input data into a latent representation, while the decoder reconstructs the original data from this representation. Attention mechanisms allow the model to selectively focus on the most relevant parts of the input data during predictions, much like humans concentrate on the most relevant details when processing complex information. Multi-head attention extends this concept by enabling the network to focus on multiple parts of the input simultaneously, capturing diverse patterns in the data. We also compare the results derived via the GMHA-AE to those derived via an analytical model that combines the shortest-path-length and search-information algorithms^10,21^, incorporating communication via both direct and indirect structural connections.

Electrophysiological FC contains rich information about the frequency characteristics of the signals in the brain, and reflects connectivity at the milli-second time scale, i.e., the actual time scale of functional interactions in the human brain. Recent work showed that the ability to predict MEG-measured FC via analytical models depends on both the frequency band of the FC and the structural measure used to assign strength to the connections of the SC matrices^17^. In this work, we test whether that is also the case when a GMHA-AE model is used to relate FC to SC.

Due to the similarities of connectivity patterns in healthy brains, several studies have used group-averaged structural and functional connectomes when calculating the SC-FC relationship. Although undoubtedly informative, those analyses cannot capture individualized aspects of the SC-FC relationship such as age dependence. A question that naturally arises is how specific to a given participant the group-derived relationship is. This is also of interest when considering populations that span a wide age range (because the SC-FC relation changes across the lifespan^22,23^) or when studying patient populations^24^.

With the above-mentioned considerations in mind, in this work we answer the following questions:

1. Is the GMHA-AE or the analytical model better at predicting electrophysiological resting-state FC from SC?
2. Does the predictive ability of each model depend on the MEG frequency band for which the resting-state FC is calculated?
3. Does the predictive ability of each model depend on the structural measure that is used to quantify connection strength in the SC matrices?
4. Is there an effect of participant age in the predictive ability of each model?
5. Is there an effect of disease in the predictive ability of each model?

## 2 Methods

The study was approved by the Cardiff University School of Psychology Ethics Committee (EC.18.08.14.5332RA3). All participants gave written informed consent. All methods were performed in accordance with the relevant guidelines and regulations.

A schematic of the analysis pipeline is shown in Fig. 1.

**Figure 1.**
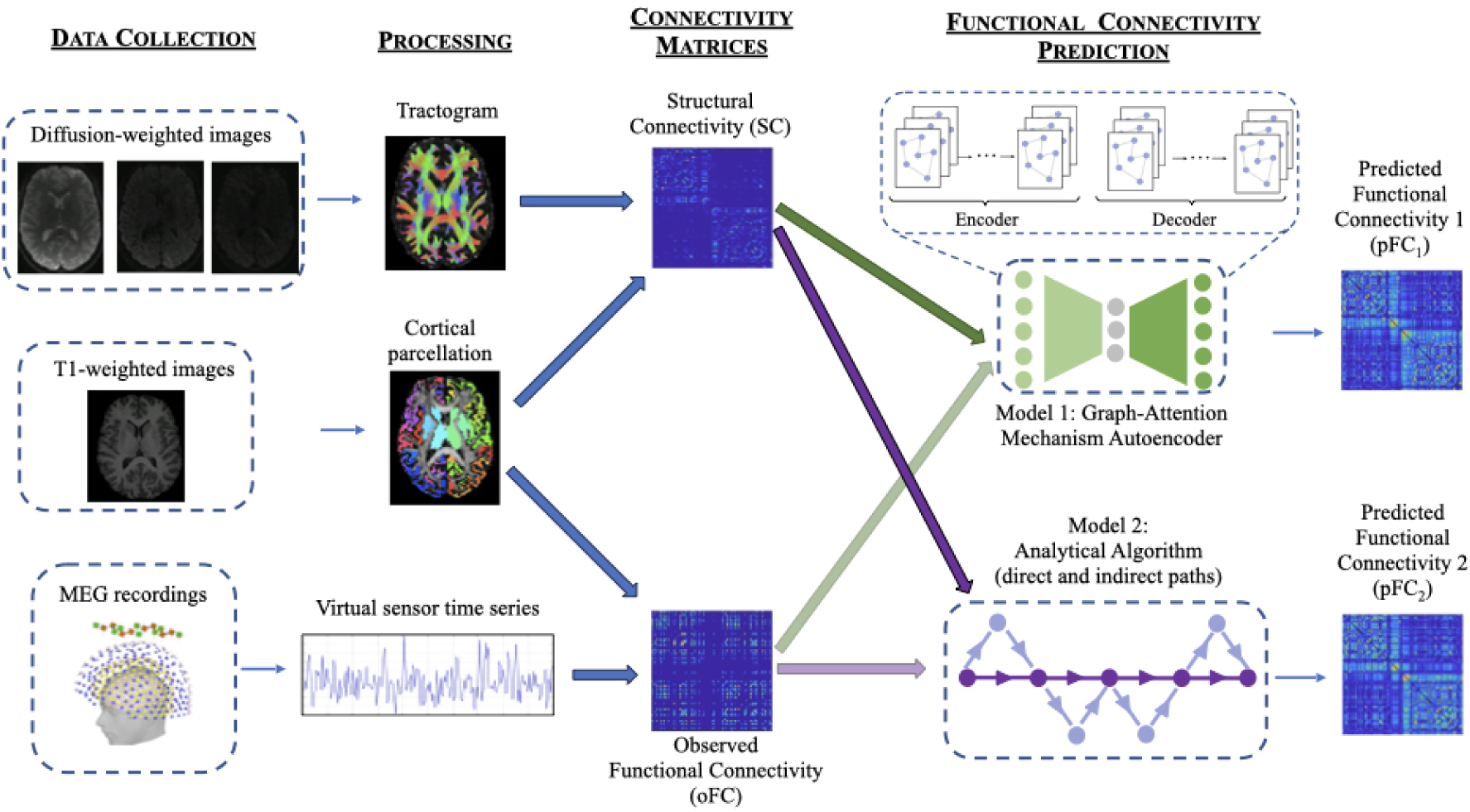
Analysis pipeline. From left to right: *Data collection* comprises MRI data collected on a 3T Connectom scanner and MEG data collected on a 275-channel gradiometer. *Processing* includes cleaning of the data, tractography, cortical parcellation, MEG source localization, and results in tractograms, cortical parcellations and activation time series for each participant. *Structural connectivity matrices* are generated from the tractograms by mapping them onto the cortical parcellations. *Functional connectivity matrices* are generated from the MEG time series via amplitude-amplitude correlations. The matrices are then fed into two distinct algorithms for *functional connectivity prediction:* the Graph-Attention Mechanism Autoencoder and the Analytical Algorithm.

### 2.1 Data Acquisition and Processing

#### 2.1.1 Sample

MRI and MEG data were collected from 126 healthy participants between the ages of 18 and 50 (73 female), 101 of whom were between the ages of 18 and 35 (55 female) via the Welsh Advanced Neuroimaging Database (WAND) study^25,26^.

Data were also collected from 5 participants with psychosis, between the ages of 18 and 35 (3 female). Four of the psychosis participants had been diagnosed between 9 and 12 months before taking part in the study, while one was diagnosed 6 years before taking part. The psychosis participants were on anti-psychotic medication. Table 1 shows the Positive and Negative Syndrome Scales (PANSS) scores^27^ for the psychosis participants, which are a measure of the severity of symptoms in the three categories of psychotic symptoms, i.e., positive, negative, and general psychopathology.

**Table 1.** PANSS scores of the psychosis participants.

| Psychosis participant | Total | Positive | Negative | General |
| --- | --- | --- | --- | --- |
| 1 | 37 | 9 | 8 | 20 |
| 2 | 59 | 11 | 18 | 30 |
| 3 | 49 | 14 | 13 | 22 |
| 4 | 41 | 9 | 9 | 23 |
| 5 | 56 | 16 | 11 | 29 |

The details of the acquisition^28^ and pre-processing are described in the Supplementary material. The pre-processing was performed using FreeSurfer^29–41^, FSL^42–44^ and MRtrix^43,43,45–58^ for the MRI data, and Fieldtrip^59^ and in-house pipelines as previously described^60^ for the MEG data.

#### 2.1.2 Connectivity Matrices

The result of this part of the analysis are connectivity matrices for each participant. The matrices are derived with the nodes being the cortical and subcortical brain areas defined via the Desikan-Killiany atlas^61^.

There are 5 SC matrices; in each the edges are weighted with a different structural measure: number of reconstructed streamlines (NS), volume-normalized NS (NS/v), fractional anisotropy (FA), inverse radial diffusivity (iRD), and total restricted signal fraction (FRt). Our choice of edge weights is motiveated by the fact that SC matrices weighted with those are good at predicting functional connectivity from both fMRI^2,10^ and MEG^17^, and provide good reproducibility for the structural connectivity matrices and their graph theoretical properties when used as edge weights^62,63^. Additionally, the FA is related to the myelination and axonal characteristics of white matter tracts and therefore is a good proxy for structural connectivity^10,17,64–66^, the iRD is related to the myelination of the white matter tracts^67–69^ and has been used in studies in health and disease^17,70^, and the FRt^71^ is attributed to water within the intra-axonal space and therefore related to axonal characteristics of the white matter tracts.

There are 4 resting-state FC matrices, one for each of the 4 MEG frequency bands: delta (1-4 Hz), theta (4-8 Hz), alpha (8-13 Hz) and beta (13-30 Hz). These 4 matrices reflect the observed FC (oFC).

Finally, there is the Euclidean distance (ED) matrix of each participant, in which each edge is the distance between the centers of the Desikan-Killiany brain areas^61^.

### 2.2 Prediction of FC

Our aim was to achieve a mapping from a participant’s SC to that participant’s FC. The resting-state FC from each frequency band was predicted for each participant individually via each of the two models (GMHA-AE and analytical), using that participant’s structural connectome, separately for each of the 5 edge weightings. Gõni et al^10^ showed that, when an analytical model is used, the prediction of functional connectivity from structural connectivity is improved by including the ED matrix as a predictor. For that reason, we include the ED matrices in the analytical model. In order to keep the GMHA-AE model comparable to the analytical one, we included the ED matrices alongside the SC matrices when predicting the FC via the GMHA-AE model, thus incorporating both connection strength (via the SC matrices) and anatomical proximity (via the ED matrices) in our models. Each of the two models resulted in 20 predicted FC (pFC) matrices for each participant, one for each frequency-band/edge-weight combination.

#### 2.2.1 GMHA-AE Model

We constructed a model based on a GMHA-AE, the structure of which is depicted in Fig. 1. The encoder, equipped with multi-head attention, processes the input data SC (which include the Euclidean distance structural matrices) to generate a latent representation that captures both direct and indirect connections within the brain. Specifially, the ED matrix is an additional input, alongside each of the SC matrices. The latent representation is then used by the decoder to reconstruct the FC matrix. By reconstructing the FC matrix, the decoder implicitly models the mapping from SC to FC, completing the autoencoder’s function. Throughout this process, the multi-head attention mechanism provides a flexible approach that captures the complex relationships inherent in brain connectivity data. More details on the autoencoder are provided in the Supplementary material, including detailed descriptions of the handling of the data in the encoder and the decoder.

We used a five-fold cross-validation methodology as follows: The sample was randomly divided into five equal sets. In each iteration, one set was selected as the validation set, and the remaining sets were used as the training set. This process continued until each set had been used as the validation set once. Parameters were optimized and selected using grid search based solely on the training data. The total number of training epochs was set to 200, with a batch size of 32, a learning rate of 1 × 10^−3^, and *γ* in the objective function set to 0.01.

The GMHA-AE model was first trained on the 126 healthy participants and used to predict their FC. For the analysis that pertains to the participants with psychosis, it was first trained on the 101 healthy participants that were 18 - 35 years old (for a detailed explanation of this choice, please see below). Once trained and validated, the data of the psychosis participants were passed through it, and their FC was predicted.

#### 2.2.2 Analytical Model

A number of algorithms that model communication between brain areas were described in an earlier publication^10^ and implemented in the Brain Connectivity Toolbox^72^. These algorithms calculate the potential predictors of FC based on a SC matrix, employing different methods to identify the optimal structural links between brain areas, and then use regression to generate a pFC that best matches the oFC. Many algorithms assume that stronger structural connections are more likely to be utilized, and several of them take into account both direct and indirect structural connections between brain areas.

In this work, we used the shortest-path-length (SPL) and search-information (SI) algorithms^10^ to derive two predictors for the analytical model. This was used for the healthy participants and the participants with psychosis. The ED matrices of the participants were used in the analytical model as an additional predictor because, as mentioned earlier, they can increase the model’s predictive ability^10,17^.

### 2.3 Statistical analysis

#### 2.3.1 Characteristics of the connectivity matrices

The 5 SC matrices derived for each participant, each weighted with a different structural measure, are intended to represent the structural connectome of that participant. To quantify how different these matrices are and evaluate whether they would result in differences when predicting FC from SC, we vectorized each of the 5 SC matrices of each participant keeping all unique connections, and concatenated the vectors for the 126 participants. This resulted in one vector for each structural measure. We then calculated the correlation coefficients between vectors for each pair of structural measures in order to quantify how correlated the matrices resulting from each structural measure are across the group of healthy participants.

#### 2.3.2 Prediction accuracy

To evaluate the accuracy of the predictions, we calculated the correlation coefficients between the oFC for each frequency band and the pFC generated for that frequency band with each of the 5 SC matrices, for the two models. To quantify how the choice of edge weight impacted the ability of the structural connectome to predict functional connectivity we used a paired t-test to compare the distributions of the oFC-pFC correlations for the 5 SC edge weights, for each frequency band, for each of the two models.

#### 2.3.3 Individuality of the SC-FC relationship

To test how "individual" (i.e., specific to each participant) the relationship between FC and SC is, we shuffled the order of the SC matrices of the 126 healthy participants while keeping the order of the FC matrices the same, and repeated the FC prediction with the two models. Thus, the FC of each healthy participant was predicted using the SC of another healthy participant. We then calculated the correlation coefficient between pFC and oFC. We repeated this process 500 times for each of the two models, and derived the mean of the correlation coefficients for each healthy participant.

#### 2.3.4 Age dependence

To evaluate the impact of participant age on the relationship between SC and FC, we calculated the correlation coefficients between the participant age and the oFC-pFC correlation coefficients, for both the GMHA-AE and the analytical model, for the 126 healthy participants.

#### 2.3.5 Psychosis

We tested the hypothesis that the SC-FC relationship is altered as a result of psychosis in comparison to healthy participants. People with psychosis exhibit brain alterations that mimic aging, such as cognitive dysfunction^73^, dendritic spine loss^74,75^ and cortical atrophy^76^. To prevent the results of our comparison being diluted by those aging-like effects, we compared the participants with psychosis to the healthy participants that matched the ages of the people with psychosis, i.e., in the age range of 18 to 35 years. This left 101 healthy participants to perform the comparison on.

To compare the connectomes of participants with psychosis to those of the healthy participants we calculated correlations between the respective connectivity matrices, for each frequency band for the FC matrices and each edge weight for the SC matrices. These correlations were then compared with the correlations of the SC matrices and the FC matrices for the healthy participants.

We used a Mann-Whitney U test to compare the distributions of the total connection strength (sum of all connections) of the healthy participants to those of the participants with psychosis, for the 5 edge weights for the SC matrices and the 4 frequency bands for the FC matrices.

To quantify any differences in the SC-FC relationship between healthy participants and participants with psychosis, we used a Mann-Whitney U test to compare the distributions of the oFC-pFC correlation coefficients between participants with psychosis and healthy participants, for each of the FC-frequency-band/SC-edge-weight combinations.

To compare the performance of the two models in predicting FC for the participants with psychosis, we performed permutation testing.

All *p*-values were corrected for multiple comparisons using the false-discovery-rate (FDR) algorithm^77,78^.

## 3 Results

### 3.1 Characteristics of the connectivity matrices

The distributions of edge weights for the SC matrices weighted with different structural metrics (Fig. 2) indicate the distinct nature of the SC networks they represent. The distributions of edge weights for the FC matrices in the 4 frequency bands are shown in (Fig. 3).

**Figure 2.**
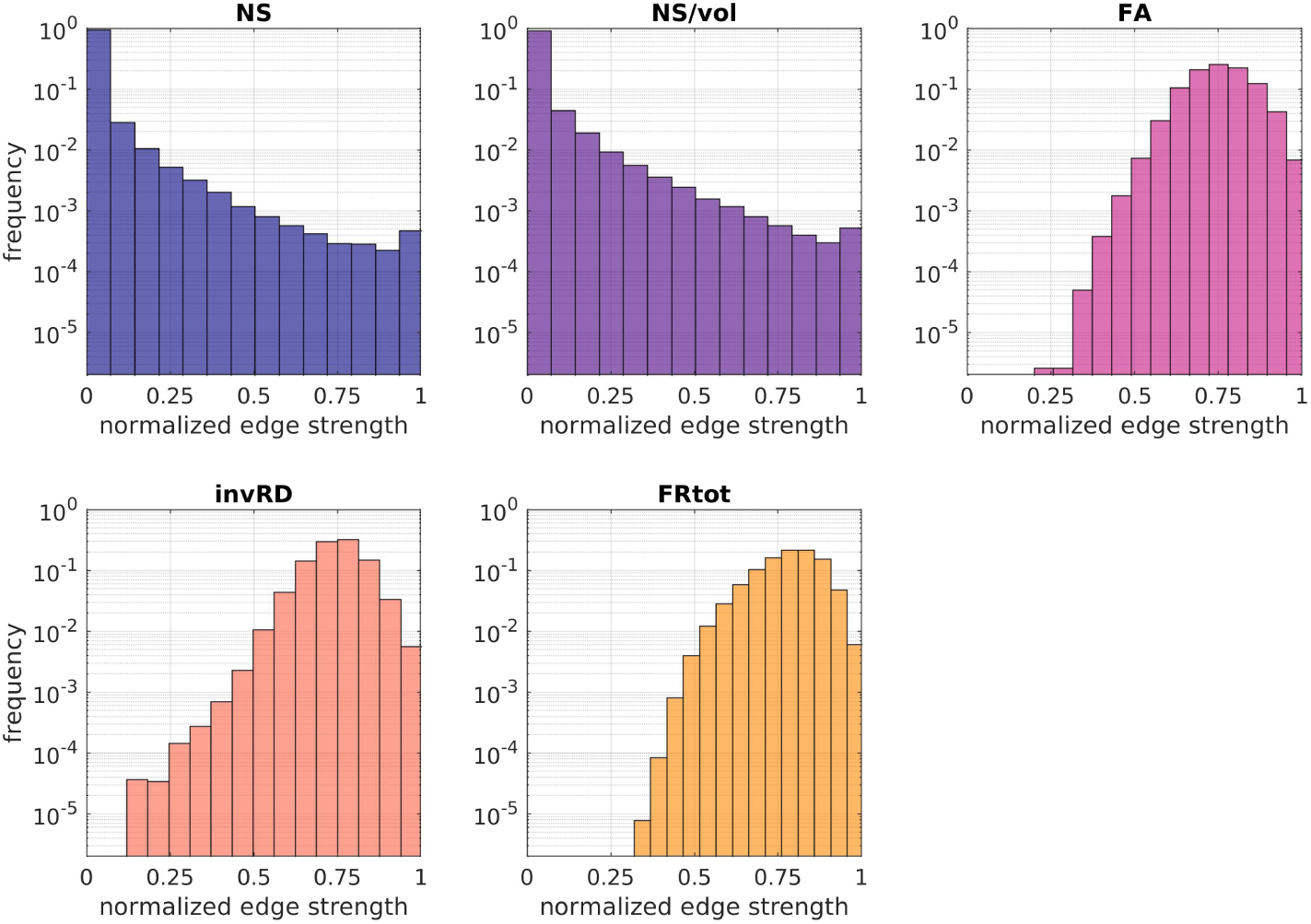
Histograms of the fraction of edge weights of the normalized SC matrices for the 5 choices of edge weight. The histograms are over all connections and all healthy participants.

**Figure 3.**
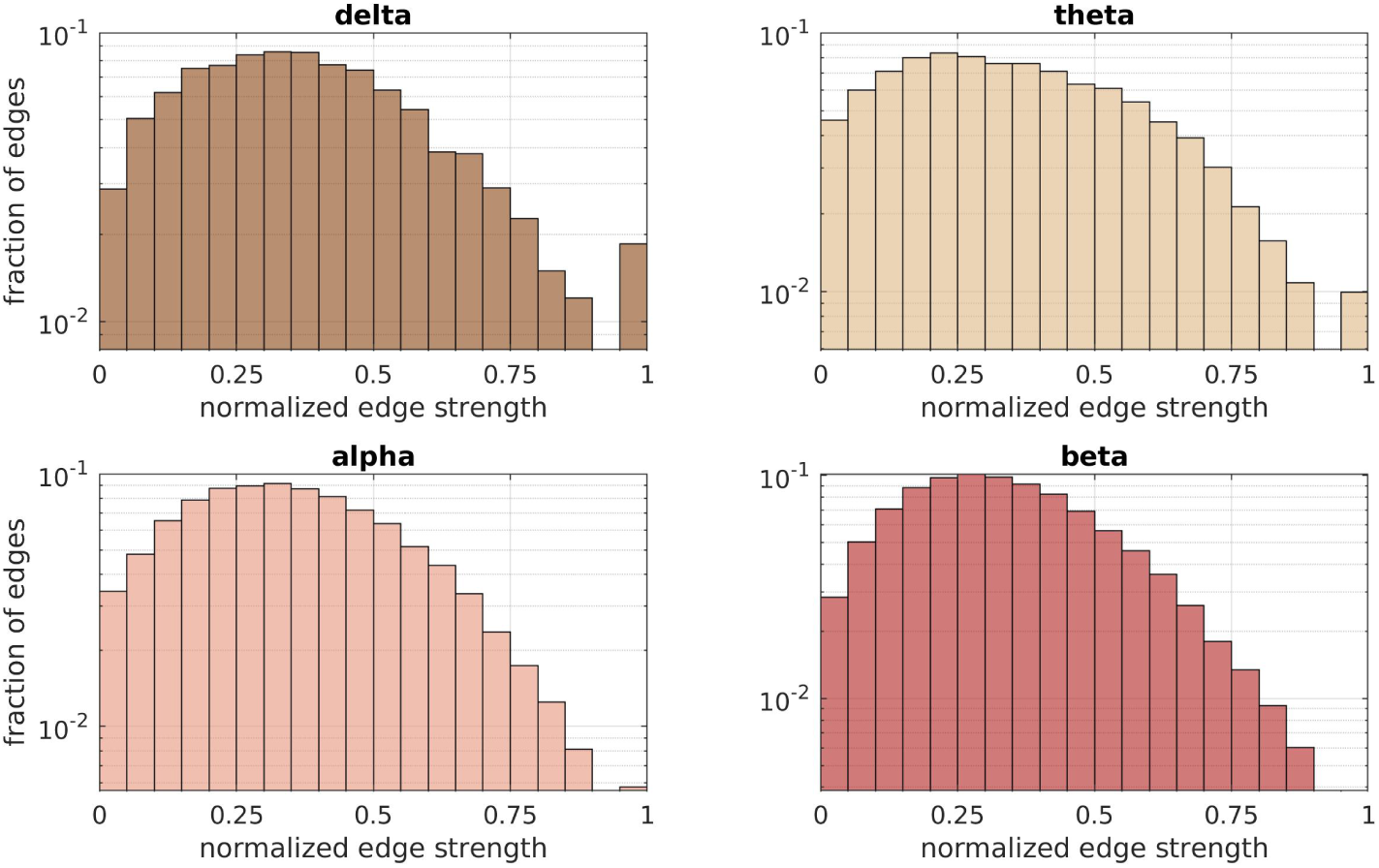
Histograms of the fraction of edge weights of the normalized FC matrices for the 4 frequency bands. The histograms are over all connections and all healthy participants.

### 3.2 Comparison of connectivity matrices of healthy participants to psychosis participants

The SC and FC matrices of the participants with psychosis were compared to the matrices of the 101 healthy participants who were between the ages of 18 and 35 years.

The NS-weighted and NS/v-weighted SC matrices of the psychosis participants exhibited high correlations with those of the healthy participants, while the ones for the other edge weights did not (left panel of Fig. 4). However, all those correlations were comparable to the correlations between SC matrices of the healthy participants (the means of the correlations for healthy participants are shown in the same figure as dashed lines for comparison).

**Figure 4.**
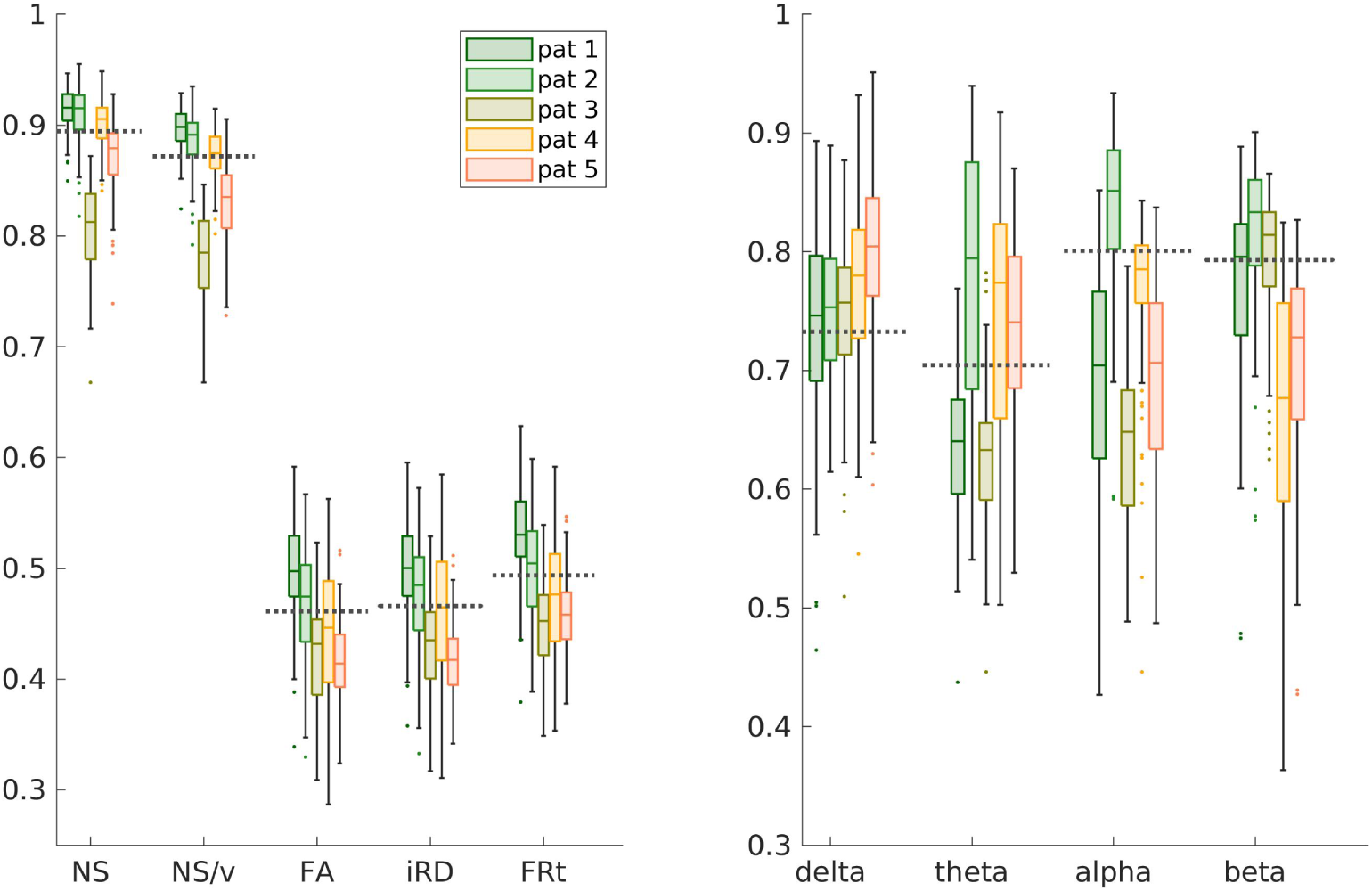
Distributions of correlations between SC (left panel) / FC (right panel) matrices of people with psychosis to those of the healthy participants, for each edge weight / frequency band. The dashed lines show the mean of the correlations between the matrices of the healthy participants for comparison. The labels ‘pat 1-5’ indicate the 5 different psychosis participants.

The FC matrices of the people with psychosis exhibited moderate correlations with the FC matrices of the 101 healthy participants aged 18 - 35 years (right panel of Fig. 4) but those correlations were also comparable with those between FC matrices of healthy participants.

We compared the total connectivity strength of the SC and FC matrices of the participants with psychosis to those of the healthy participants (Fig. 5). The *p*-values resulting from the Mann-Whitney U tests between the distributions of the connectivity strength for the healthy participants and those with psychosis are given in that figure. The only *p*-value that was under 0.05 is the one for the beta band FC, but it did not survive FDR multiple-comparison correction.

**Figure 5.**
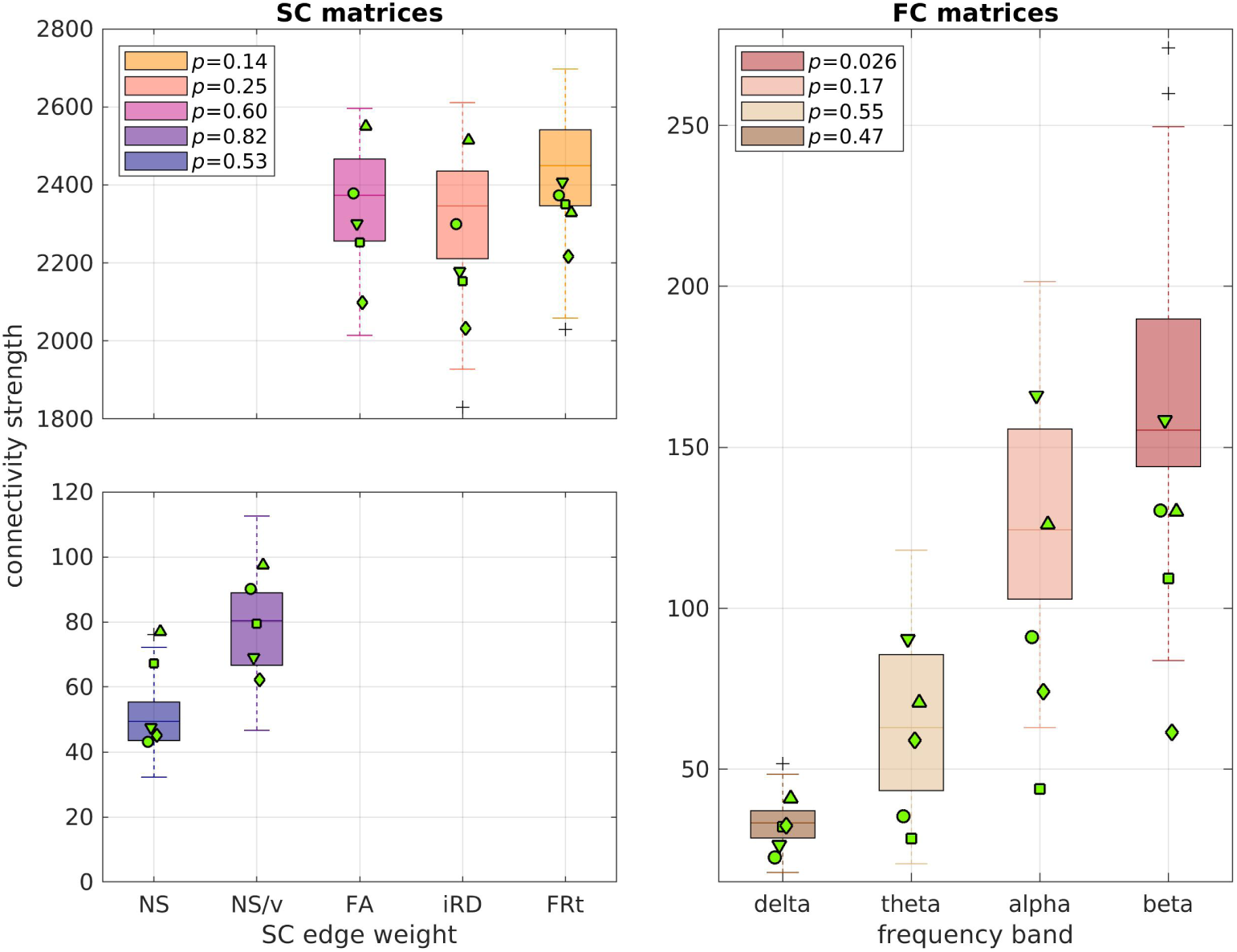
Distributions of the connection strength of the healthy participants (boxplots) in comparison to the connection strength of the psychosis participants (green markers). Each marker indicates one of the 5 participants with psychosis. The *p*-values of the the Mann-Whitney U test for the total edge strength distributions between healthy participants and those with psychosis are given in the legend. None of the values survived FDR multiple-comparison correction.

Our results for the SC matrices are in agreement with those of earlier work^79^, which demonstrated no differences in the NS-based total SC strength and a small reduction in the FA-based total SC strength (in our cohort, the latter showing a trend towards smaller values for the psychosis participants in comparison to the healthy participants; respective means: 2300 and 2370). They are also in agreement with the results reported in a different study^80^, considering the fact that most of our psychosis participants had been recently diagnosed.

Our results for the FC matrices are in agreement with the results presented previously^81^, where no statistically significant differences were found between the EEG amplitude-amplitude connectivity of schizophrenia patients and healthy participants. As pointed out in the review article^82^, there is heterogeneity in the results coming from MEG- and EEG-based connectivity studies of psychosis.

### 3.3 Prediction of FC with the GMHA-AE model

#### 3.3.1 Maintaining the participant link between SC and FC

The GMHA-AE model was good at predicting the oFC (left panel of Fig. 6) with the mean of the correlations being above 0.8 for the alpha and beta frequency bands and above 0.4 for the delta and theta bands. The only exception was the prediction of the oFC with the NS-based SC matrices for the delta band, for which the correlation coefficients between oFC and pFC were low.

**Figure 6.**
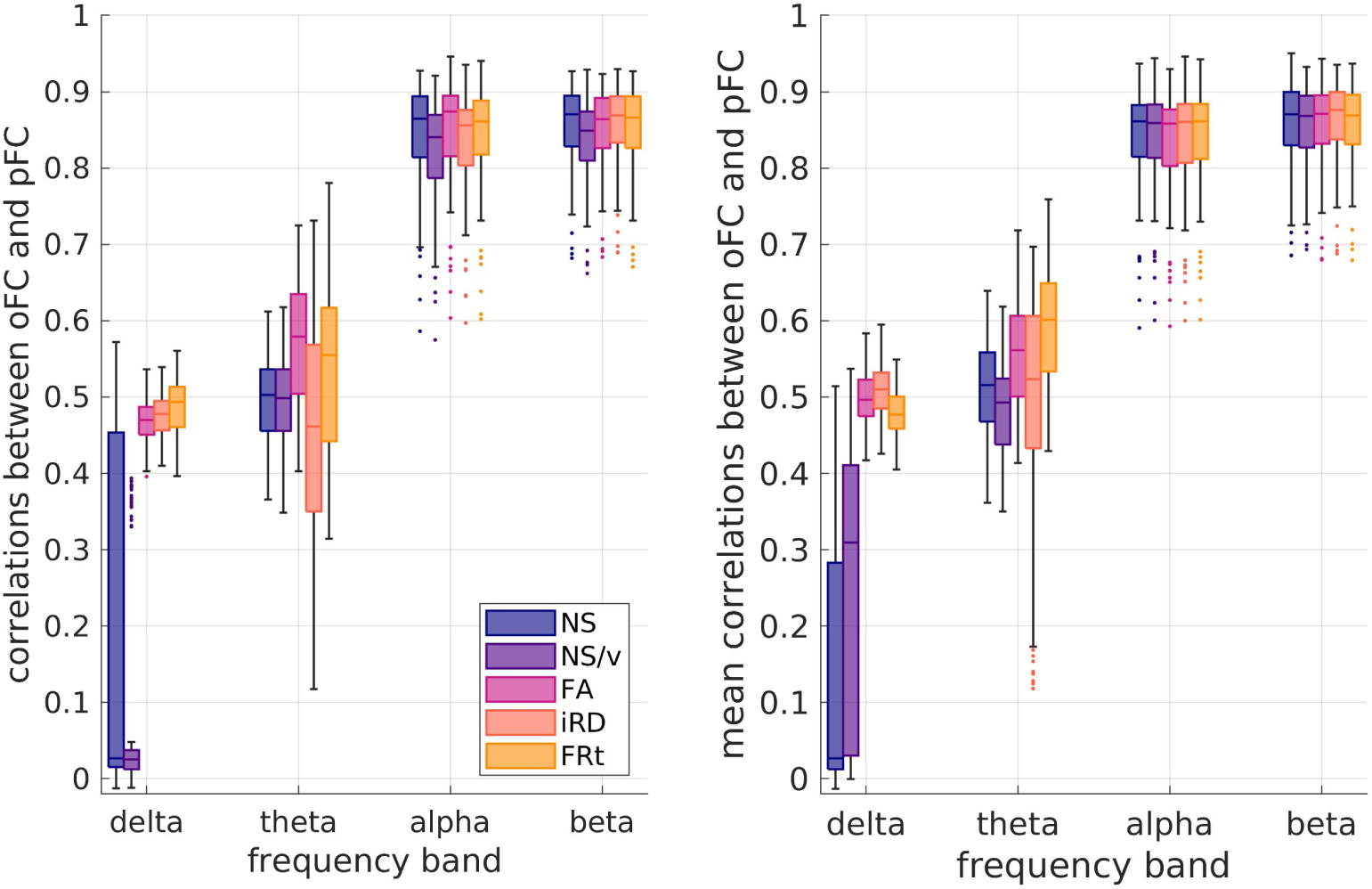
Left panel: Distributions (over the 126 participants) of the correlation coefficients between oFC and pFC derived with the GMHA-AE model, when no permutations of the SC matrices have been done. Right panel: Distribution (over the 126 participants) of the mean (over the 500 shufflings) of the oFC-pFC correlations derived with the GMHA-AE model.

#### 3.3.2 Breaking the participant link between SC and FC

The pFC calculated via the GMHA-AE model after shuffling the SC matrices of the healthy participants exhibited a level of correlation with the oFC similar to that when the SC matrices of the participants had not been shuffled (right panel of Fig. 6). We note that the figures on the two panels of Fig. 6 are not directly comparable, because the left panel shows the distribution of correlations over the 126 participants when the SC matrices have not been shuffled, while the right panel shows the distribution over the 126 participants of *the mean* over the 500 shufflings of the oFC-pFC correlations.

#### 3.3.3 Age dependence

For the pFC calculated via the GMHA-AE model, we found no statistically significant correlations between participant age and the oFC-pFC correlation coefficients after FDR correction (Fig. 7). This indicates that the predictive ability of the GMHA-AE model is stable across ages and not altered with participant age, at least for healthy adults in the age range of 18 - 50 years.

**Figure 7.**
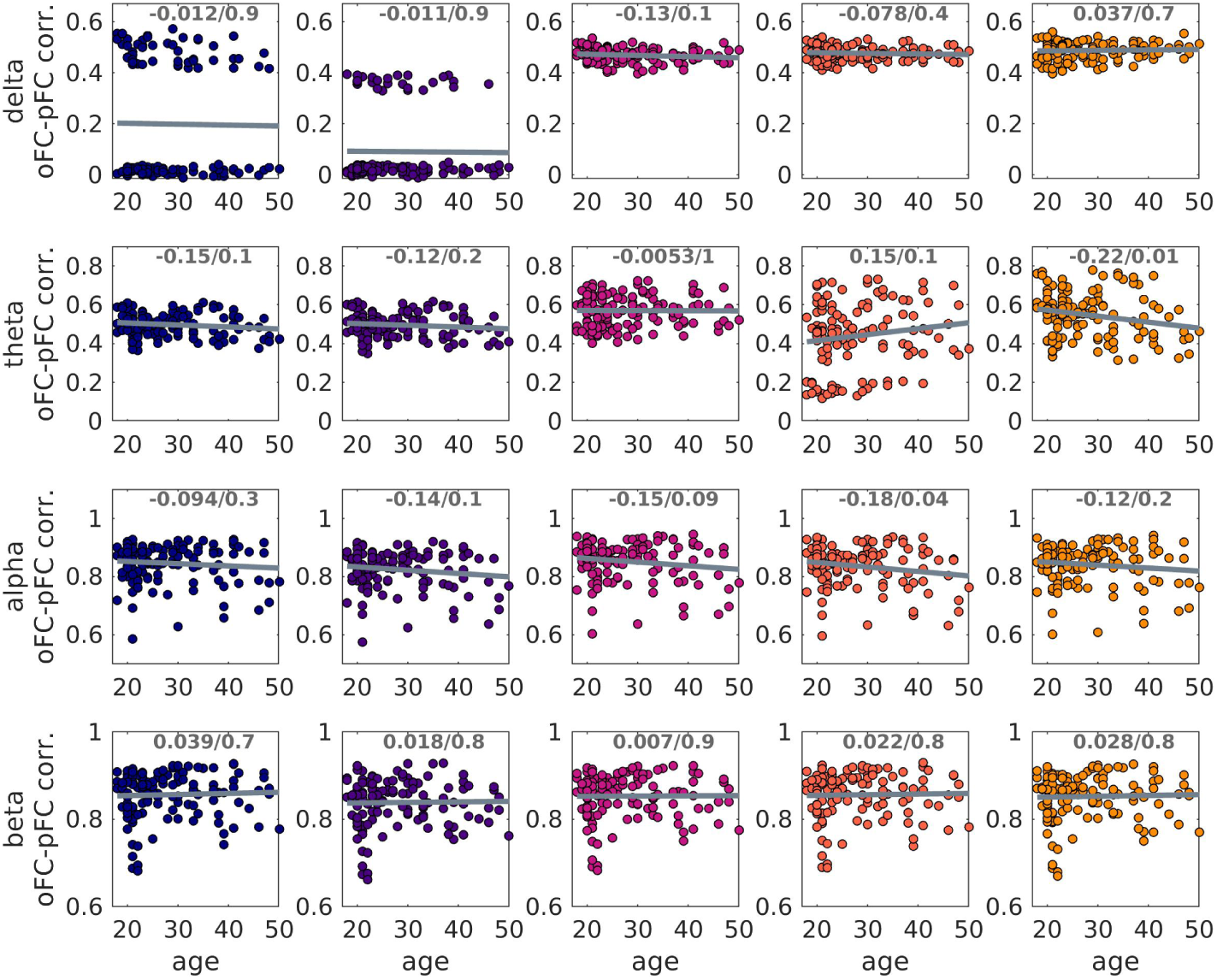
Scatter plots of the oFC-pFC correlation coefficients versus age. The 5 columns refer to the 5 SC edge weights, from left to right: NS, NS/v, FA, iRD and FRt. The best-fit line is shown. The correlation coefficients and *p*-values are also shown at the top of each plot. None of the *p*-values were statistically significant after FDR correction.

#### 3.3.4 Psychosis

When the GMHA-AE model that had been trained on healthy participants was used to calculate the pFC of the psychosis participants, the correlations between oFC and pFC were lower for all frequency bands and all SC edge weights compared to the healthy participants (Fig. 8). The *p*-values of the Mann-Whitney U test comparing the distributions of the healthy participants to those with psychosis were all less than 2 ×10^−4^ (Table 3) and survived FDR correction, with the exception of the NS/delta and FA/theta pairs (*p* = 0.48 and 0.10 respectively).

**Figure 8.**
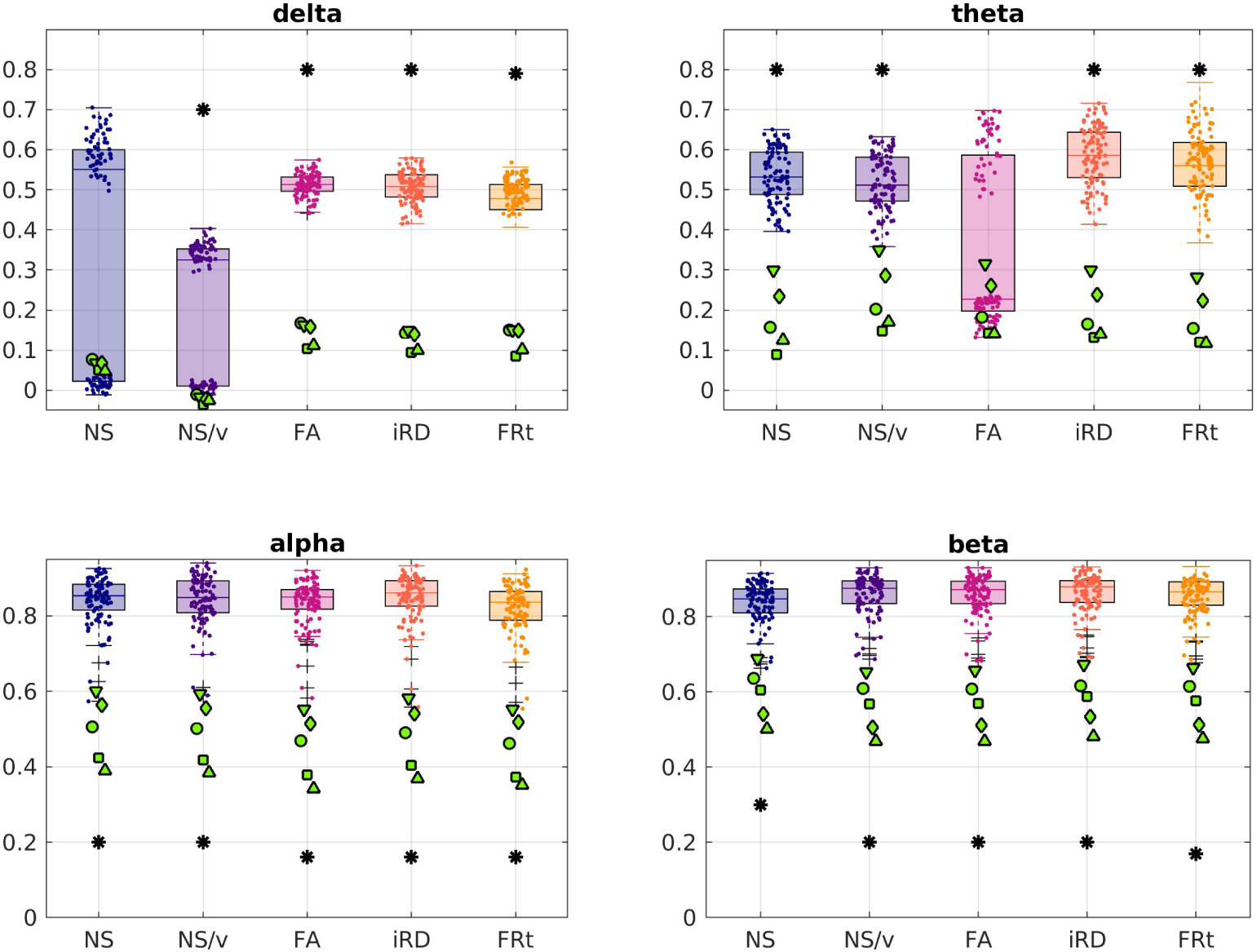
The oFC-pFC correlations of the psychosis participants (green markers) overlaid on the corresponding correlation distributions of the healthy participants (boxplots and clouds of points), for the case in which the FC is predicted with the GMHA-AE model. Each marker shape indicates a different psychosis participant. Asterisks indicate statistically significant differences (after FDR correction) between the distributions for the healthy participants and those with psychosis.

### 3.4 Prediction of FC with the analytical model

#### 3.4.1 Maintaining the participant link between SC and FC

When the analytical model that combines the SPL and SI algorithms was used to predict the healthy participants’ FC, the correlations between the oFC and pFC depended on the frequency band of the observed FC and on the edge weight used in the SC matrices, as shown on the left panel of Fig. 9. The higher frequency bands resulted in higher correlations between observed and predicted FC, indicating that a combination of the SPL and SI algorithms is better at predicting FC in higher rather than lower frequencies. Additionally, NS-based structural metrics (NS and NS/v) were worse at predicting FC in all 4 frequency bands considered as opposed to the other 3 structural metrics.

**Figure 9.**
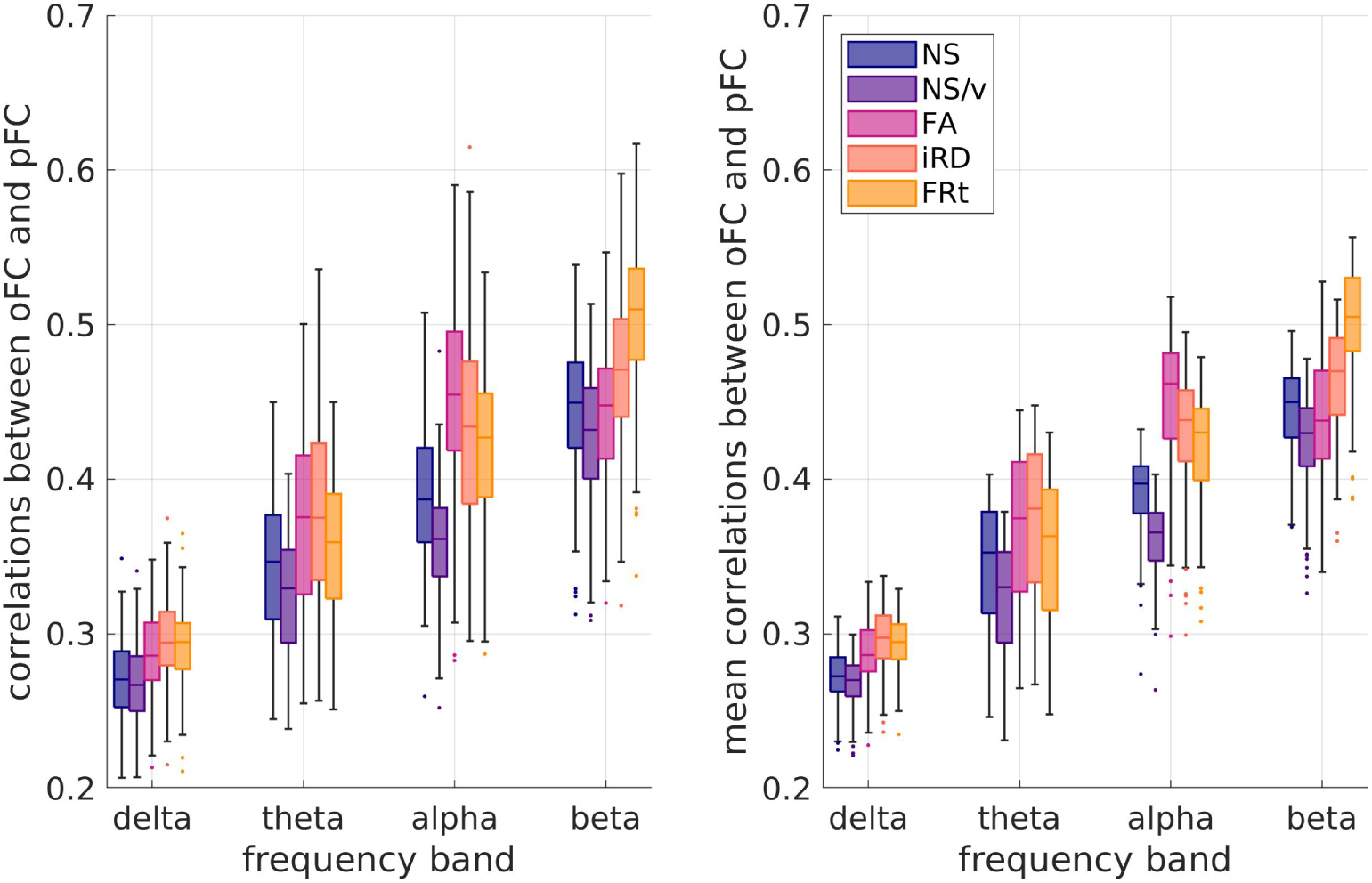
Left panel: Distributions (over the 126 participants) of the correlation coefficients between oFC and pFC derived with the analytical model when no permutations of the SC matrices have been done. Right panel: Distributions (over the 126 participants) of the mean values (over permutations) of the oFC-pFC correlations derived with the analytical model.

The paired t-test between the correlation distributions for each pair of SC edge weights showed that the distributions are statistically significantly different after FDR correction with very few exceptions (Table 2). The NS-based SCs were generally worse at predicting the oFC than the other 4 edge weights (Fig. 9).

**Table 2.** *p*-values for the t-test between the correlation distributions for oFC and pFC for pairs of SC edge weights for each frequency band. Left 4 columns: prediction with the autoencoder model; right 4 columns: predictions with the analytical model. Bold-faced p-values are **not statistically significant** after FDR correction. The *p*-value threshold resulting from the application of the FDR correction was 0.003.

|  | AUTOENCODER MODEL |  |  |  | ANALYTICAL MODEL |  |  |  |
| --- | --- | --- | --- | --- | --- | --- | --- | --- |
|  | delta | theta | alpha | beta | delta | theta | alpha | beta |
| NS - NS/v | $2 \times 10^{-4}$ | **0.12** | $< 10^{-10}$ | $< 10^{-10}$ | $8 \times 10^{-3}$ | $< 10^{-10}$ | $< 10^{-10}$ | $< 10^{-10}$ |
| NS - FA | $< 10^{-10}$ | $< 10^{-10}$ | $10^{-4}$ | $5 \times 10^{-5}$ | $< 10^{-10}$ | $< 10^{-10}$ | $< 10^{-10}$ | **0.73** |
| NS - iRD | $< 10^{-10}$ | $3 \times 10^{-4}$ | $6 \times 10^{-5}$ | **0.71** | $< 10^{-10}$ | $< 10^{-10}$ | $10^{-8}$ | $3 \times 10^{-6}$ |
| NS - FRt | $< 10^{-10}$ | $4 \times 10^{-9}$ | **0.02** | $3 \times 10^{-6}$ | $< 10^{-10}$ | $4 \times 10^{-3}$ | $10^{-7}$ | $< 10^{-10}$ |
| NS/v - FA | $< 10^{-10}$ | $< 10^{-10}$ | $< 10^{-10}$ | $< 10^{-10}$ | $< 10^{-10}$ | $< 10^{-10}$ | $< 10^{-10}$ | $3 \times 10^{-3}$ |
| NS/v - iRD | $< 10^{-10}$ | $4 \times 10^{-4}$ | $< 10^{-10}$ | $< 10^{-10}$ | $< 10^{-10}$ | $< 10^{-10}$ | $< 10^{-10}$ | $< 10^{-10}$ |
| NS/v - FRt | $< 10^{-10}$ | $4 \times 10^{-9}$ | $10^{-10}$ | $< 10^{-10}$ | $< 10^{-10}$ | $< 10^{-10}$ | $< 10^{-10}$ | $< 10^{-10}$ |
| FA - iRD | 0.004 | $< 10^{-10}$ | $< 10^{-10}$ | $6 \times 10^{-4}$ | $< 10^{-10}$ | $3 \times 10^{-3}$ | $10^{-8}$ | $< 10^{-10}$ |
| FA - FRt | $6 \times 10^{-8}$ | **0.07** | $< 10^{-10}$ | **0.47** | $8 \times 10^{-4}$ | $< 10^{-10}$ | $< 10^{-10}$ | $< 10^{-10}$ |
| iRD - FRt | $10^{-9}$ | $4 \times 10^{-6}$ | **0.02** | $3 \times 10^{-3}$ | **0.05** | $3 \times 10^{-3}$ | $10^{-3}$ | $< 10^{-10}$ |

#### 3.4.2 Breaking the participant link between SC and FC

When the SC matrices of the participants were shuffled, so that the SC matrix used in the analytical model to predict the FC no longer corresponded to the same participant, small differences were observed in the correlation coefficients between oFC and pFC (Fig. 9). As previously, we note that the figures on the two panels of Fig. 9 are not directly comparable, because the left panel shows the distribution of correlations over the 126 participants when the SC matrices have not been shuffled, while the right panel shows the distribution over the 126 participants of *the mean* over the 500 shufflings of the oFC-pFC correlations.

#### 3.4.3 Age dependence

We found statistically significant correlations (after FDR multiple comparison correction) between the participant age and the oFC-pFC correlation coefficients derived with the analytical model for the iRD SCs (all frequency bands), for the FA SC matrices (theta, alpha and beta frequency bands), and the FRt SC matrices (alpha and beta frequency bands) as shown in Fig. 10. The correlations were negative, indicating a weakening of the predictive power of the analytical model as participants age.

**Figure 10.**
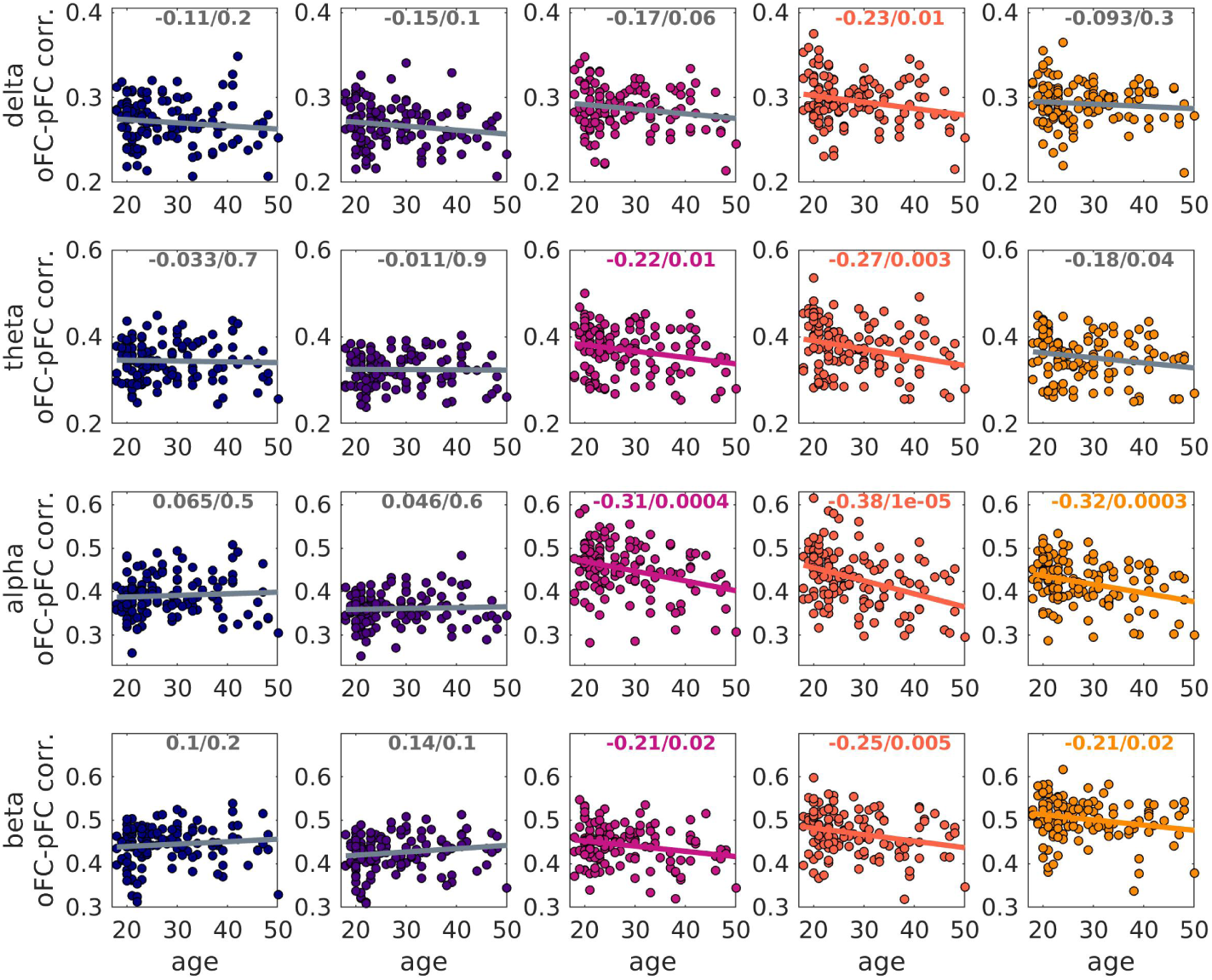
Scatter plots of the oFC-pFC correlation coefficients versus age, when the FC is predicted via the analytical model. The 5 columns refer to the 5 SC edge weights, from left to right: NS, NS/v, FA, iRD and FRt. The best-fit line is shown. The correlation coefficients and p-values are also shown at the top of each plot. Colored font / line indicate that the related *p*-value survived FDR correction, while grey indicates that it did not.

#### 3.4.4 Psychosis

A Mann-Whitney U test between homologous distributions of oFC-pFC correlations showed that the psychosis participants exhibited statistically significant differences in the correlations between oFC and pFC for the delta band (all edge weights), the alpha band (for the FA- and iRD-weighted SC matrices) and the beta band (for SC matrices weighted with FA and FRt) in comparison to the healthy participants. The correlations for the psychosis participants were higher in the delta band and lower in the alpha and beta bands (Fig. 11). This indicates that the analytical model is a better representation of the communication mechanisms between brain areas in psychosis participants than it is in healthy participants in the delta band, while the opposite is true in the alpha and beta bands.

**Figure 11.**
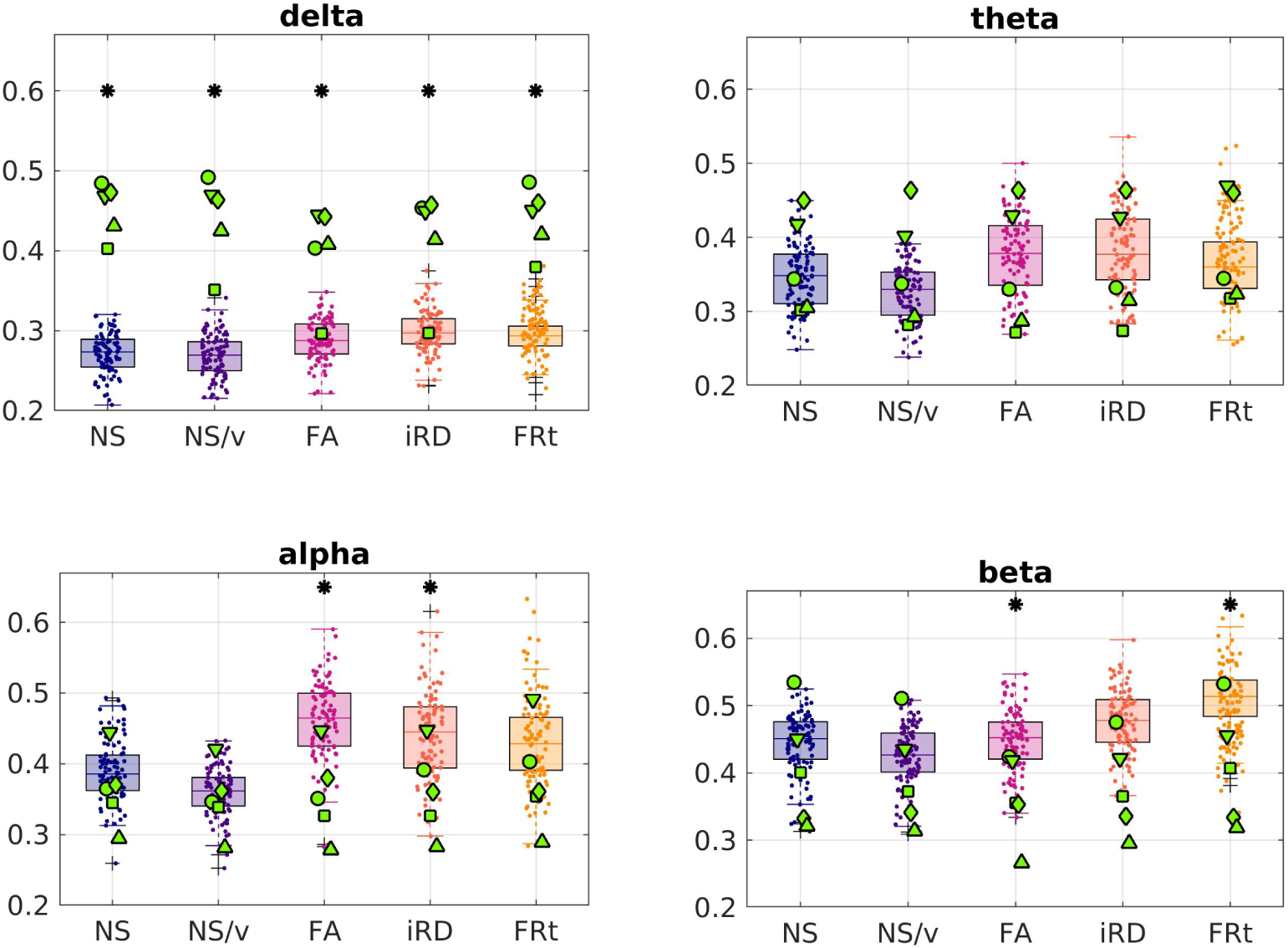
The oFC-pFC correlations of the psychosis participants (green markers) are overlaid on the equivalent correlation distributions of the healthy participants (boxplots and clouds of points), when the analytical model is used to predict the FC. Each marker shape indicates a different psychosis participant. The asterisks indicate the frequency bands and edge weights for which the two distributions are statistically significantly different after multiple comparison correction.

Table 3 gives the *p*-values for the Mann-Whitney U test between the healthy-participant and the psychosis-participant distributions of the correlations, for each frequency band and edge weight.

**Table 3.**
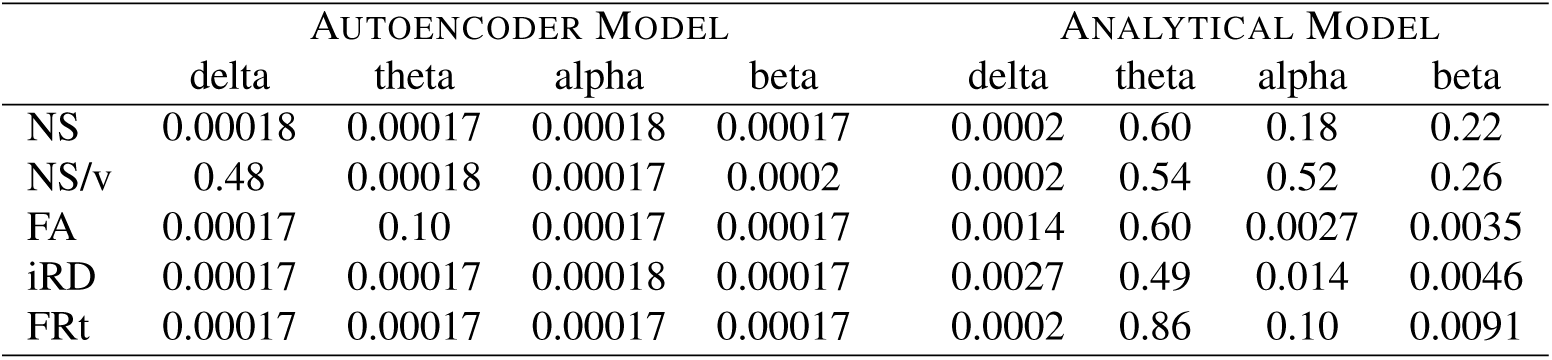
*p*-values for the Mann-Whitney U test comparing the correlation distributions for the healthy participants and the psychosis participants, when FC is predicted via the autoencoder model (left 4 columns) and the analytical model (right 4 columns). All *p*-values shown in the table that are lower than 0.05 remained statistically significant after FDR correction. The *p*-value threshold that resulted from the application of the FDR correction was 0.014.

### 3.5 Comparison of results of the two models for the psychosis participants

Fig. 12 shows the oFC-pFC correlations for the psychosis participants for the GMHA-AE and the analytical model. The distributions of the correlations are statistically significantly different for all FC frequency bands and SC edge weights. The *p*-values resulting from the permutation testing are shown in Table 4. Notably, the GMHA-AE model performs better than the analytical one for the alpha and beta frequency bands, while the opposite is true for the delta and theta bands.

**Figure 12.**
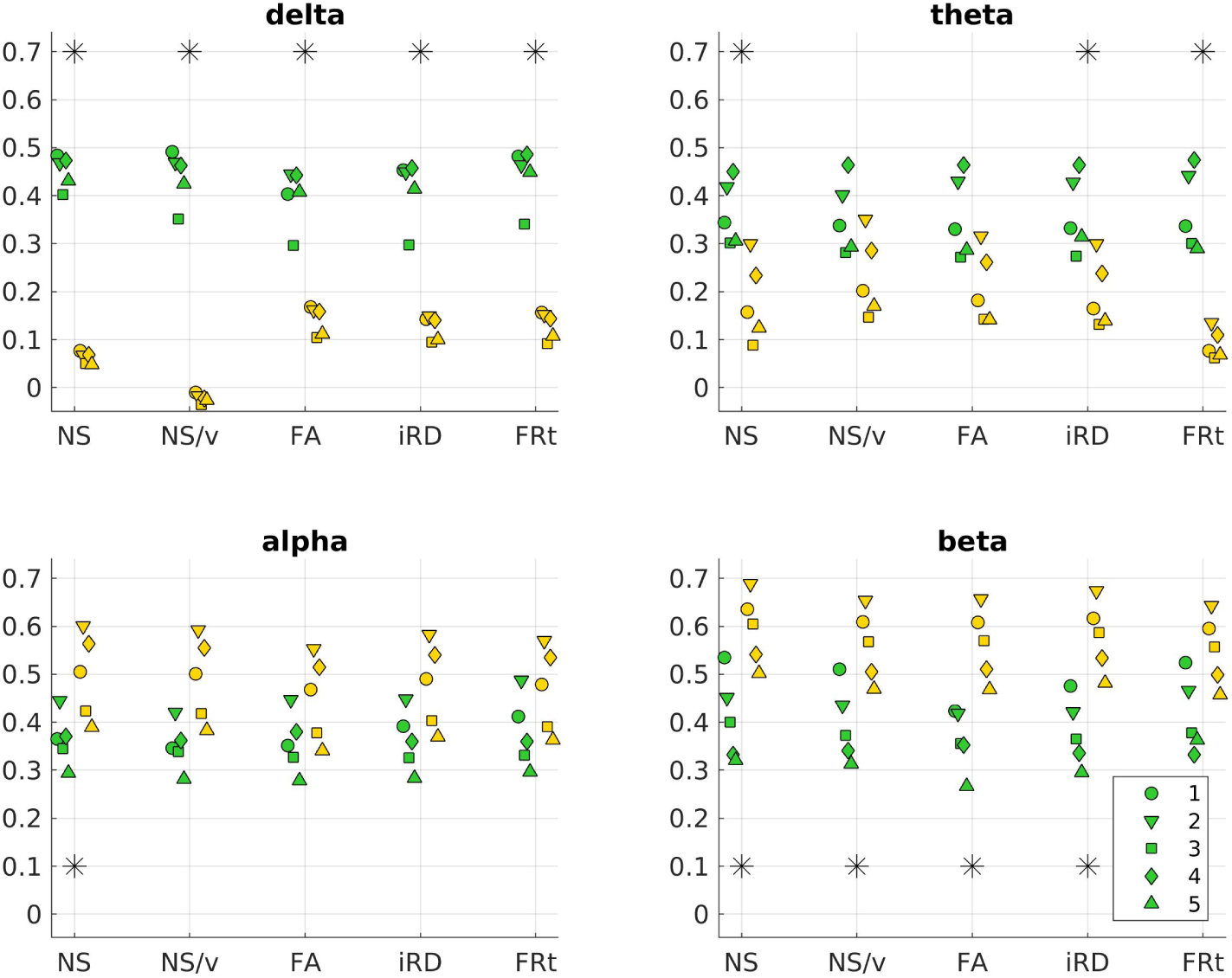
Correlations between oFC and pFC for the 5 psychosis participants for the GMHA-AE model (yellow markers) and the analytical model (green markers) across FC frequency bands and SC edge weights. Each marker shape denotes a different psychosis participant. The asterisks denote the cases for which the differences in the distributions are statistically significantly different after FDR correction.

**Table 4.** *p*-values resulting from the permutation testing to compare the correlation distributions for the psychosis participants between the GMHA-AE and the analytical model. *p*-values that remain statistically significant after FDR correction are bold-faced. The *p*-value threshold that resulted from the application of the FDR procedure was 0.012.

|  | delta | theta | alpha | beta |
| --- | --- | --- | --- | --- |
| NS | **0.004** | **0.012** | **0.016** | **0.012** |
| NS/v | **0.008** | 0.060 | 0.040 | **0.016** |
| FA | **0.004** | 0.040 | 0.096 | **0.004** |
| iRD | **0.004** | **0.012** | 0.056 | **0.012** |
| FRt | **0.004** | **0.004** | 0.127 | 0.060 |

## 4 Discussion

To the best of our knowledge, this is the first study to investigate the relationship between SC and electrophysiological resting-state FC with an attention-based deep-learning model, in both health and in psychosis.

The GMHA-AE model resulted in pFC that better matches the oFC compared to the analytical model, across the 4 frequency bands and 5 edge weights, with the exception of the delta band/NS-based edge weights combination. Additionally, in the alpha and beta frequency bands the mean of the correlation distributions was above 0.8 across SC edge weights, which is higher than the 0.55 for the distributions of the individual-level correlations reported in the fMRI-based study described in an earlier study by Sarwar et al^18^ (in which the NS was used to assign SC strength between regions). This difference in performance could be due to differences in our deep-learning model compared to that in their study^18^, the fact that we use electrophysiological FC which is not confounded by the haemodynamic response delay, or the fact that in that study^18^ they did not use information on the Euclidean distance between nodes of the networks, which can improve the FC prediction^10^. An exact comparison between our MEG-based study and their fMRI-based one is not feasible due to the different aspects of FC that the two modalities measure. When comparing our results to those of that study^18^, we do so using our results for the alpha and beta bands because those give functional connectivity that is the most comparable to fMRI-measured functional connectivity^12^. However, in accordance with the conclusion reached in that study^18^ about fMRI-measured FC, our results imply that FC measured via electrophysiological recordings is tightly linked to SC derived via tractography, and that relationship is present at individual-participant level. Therefore, existing models of communication between brain areas can be improved to incorporate sources of that interdependence^1^ that could increase the correlations.

Both the GMHA-AE and the analytical model were better at predicting the FC in the alpha and beta frequency bands in comparison to the delta and theta bands. This is in agreement with the results presented in earlier work^17^, which examined a cohort of 90 healthy participants and investigated different microstructural measures as edge weights for the SC matrices (NS, FA, iRD). That study found lower oFC-pFC correlations compared to the correlations we observed here, across functional frequency bands and SC edge weights, which could be attributed to differences in the tractography algorithm used, and the quality of the diffusion-MRI data acquired (the MRI data for this study were collected on a Connectom scanner that has high diffusion gradients as opposed to the MRI scanner used for the data collected in^17^; MEG data were collected in the same system for both studies).

The structural measure that was used as edge weight in the SC matrices impacted how well the oFC was predicted, mainly for the analytical model. Edge weights that are related to myelin or axonal characteristics (FA, iRD and FRt) resulted in better FC prediction than those related to NS when the analytical model was used, with the differences in the respective distributions being highly statistically significant (Table 2). This indicates that, if the analytical model is representative of communication between brain areas, resting-state FC is supported by myelin and axonal characteristics more than it is by the relative number of axonal projections between brain areas. When the GMHA-AE model was used, SC matrices with edge weights related to myelin and axonal measures gave better prediction for the delta and theta frequency bands (except when theta-band FC was predicted using the iRD-weighted SC matrices). That model gave a much more stable performance across SC edge weights for the alpha and beta bands, even though the distributions were statistically significantly different. This likely points to an ability of the GMHA-AE model to capture patterns in the SC matrices that pertain to communication between brain areas, regardless of the edge weights used.

We also note that for the case of the GMHA-AE model, when the delta band FC was predicted with the NS-weighted or the NS/v-weighted SC matrices, there was a subset of the sample for which the oFC-PFC correlations were high, and another subset for which the correlations were low (Fig. 6). The poor performance for the latter subset points to the fact that some of the test sets may contain individuals whose brain connectivity differs significantly from those in the training set, or that the model is learning specific characteristics from the training data that do not apply to the test data. This can make it difficult for the model to generalize properly and thus it does not perform well across all folds. This issue is present in the delta band - we observed that there is higher variability in the edge strengths in participants in the delta band compared to the other 3 bands (Fig. 13). Specifically, the median values of the variability were 0.21, 0.19, 0.18 and 0.17 for the delta, theta, alpha and beta bands respectively. The distribution of the delta band was statistically significantly different from the other 3, with *p*-values of the related t-test less than 10^−10^ in all cases. This implies that there are more varied edge strengths that the model is trying to predict in the delta band, leading to the difference in performance between the delta and the other bands.

**Figure 13.**
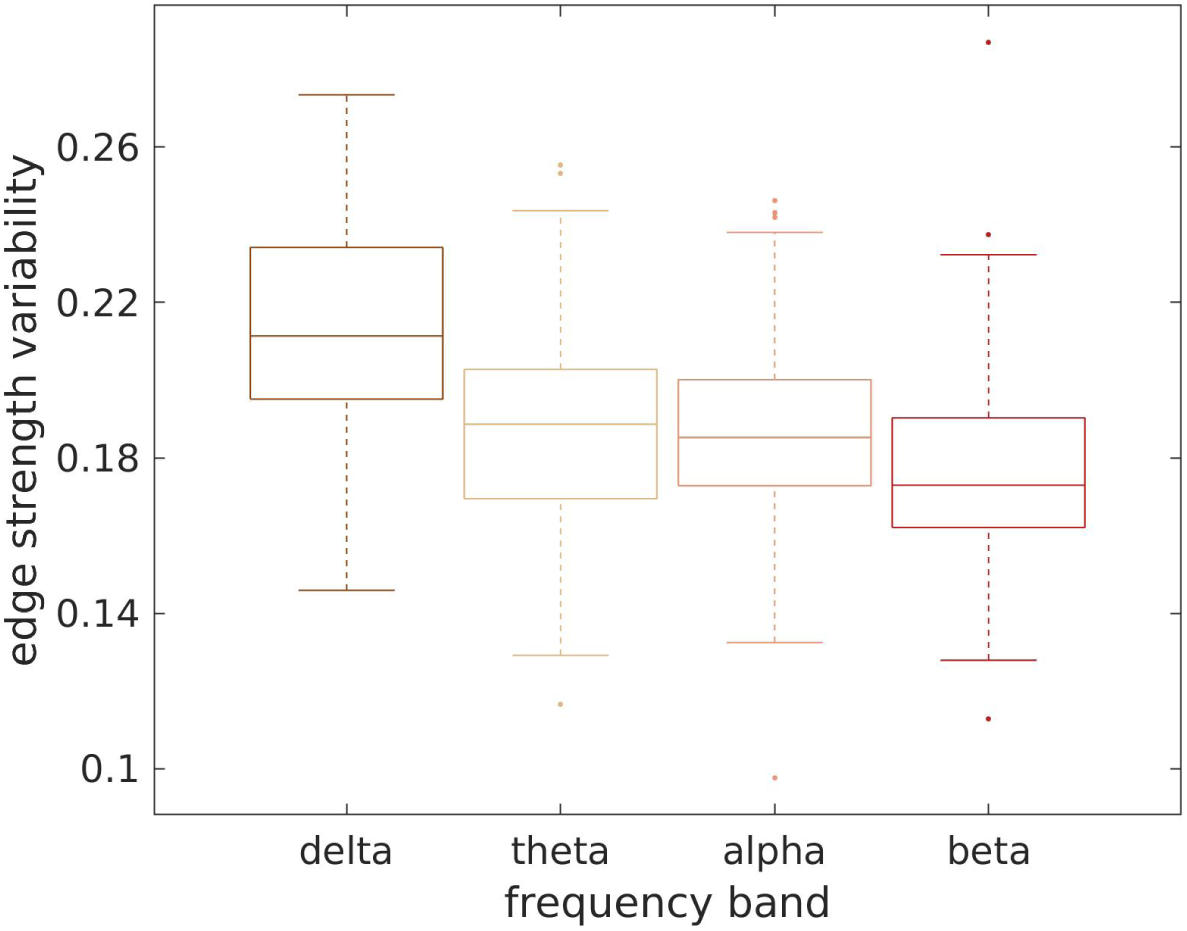
Edge strength variability in the healthy participants for the 4 frequency bands. Notably, the variability is higher in the delta band.

We observed no significant change in the predictive ability of the two models when the SC matrices of the healthy participants were shuffled (i.e., the FC of a participant was predicted by the SC of a different participant). This was true across structural edge weights and FC frequency bands. It is an indication of the similarity of the connectomes and the communication mechanisms in healthy participants, and is in contrast to the results obtained when the structural and functional matrices of the psychosis patients were used.

In order to investigate the interpretability of our GMHA-AE model and understand what features it prioritizes, we conducted an analysis on its learned latent space. We performed t-distributed Stochastic Neighbor Embadding (t-SNE) to visualize the high-dimensional latent representations of all test samples in a 2-dimensional space. The results of the analysis are shown in Fig. 14. Each point in the plot corresponds to a participants, and it is colored by the model’s prediction error (MSE) for that participant. The resulting visualization provides compelling evidence of a well-structured latent space. A distinct cluster of participants with high prediction error is clearly separated in the bottom-right corner. This group represents less typical or harder to predict participants. Conversely, the majority of participants, for which the model achieved low-to-moderate prediction error, form another large, distinct group. This structural separation demonstrates that our model has successfully learned to distinguish between typical and atypical patterns in the SC-to-FC mapping. It prioritizes features that are indicative of this predictability, effectively translating the difficulty of the prediction task into geometric separation in the latent space. We note here that a more detailed interpretability investigattion could include other methods, for example an ablation analysis or other. This falls beyond the scope of this work, which was mainly to compare the GMHA-AE model to an analytical one and to explore the impact of psychosis participants on them, and will be explored in future work.

**Figure 14.**
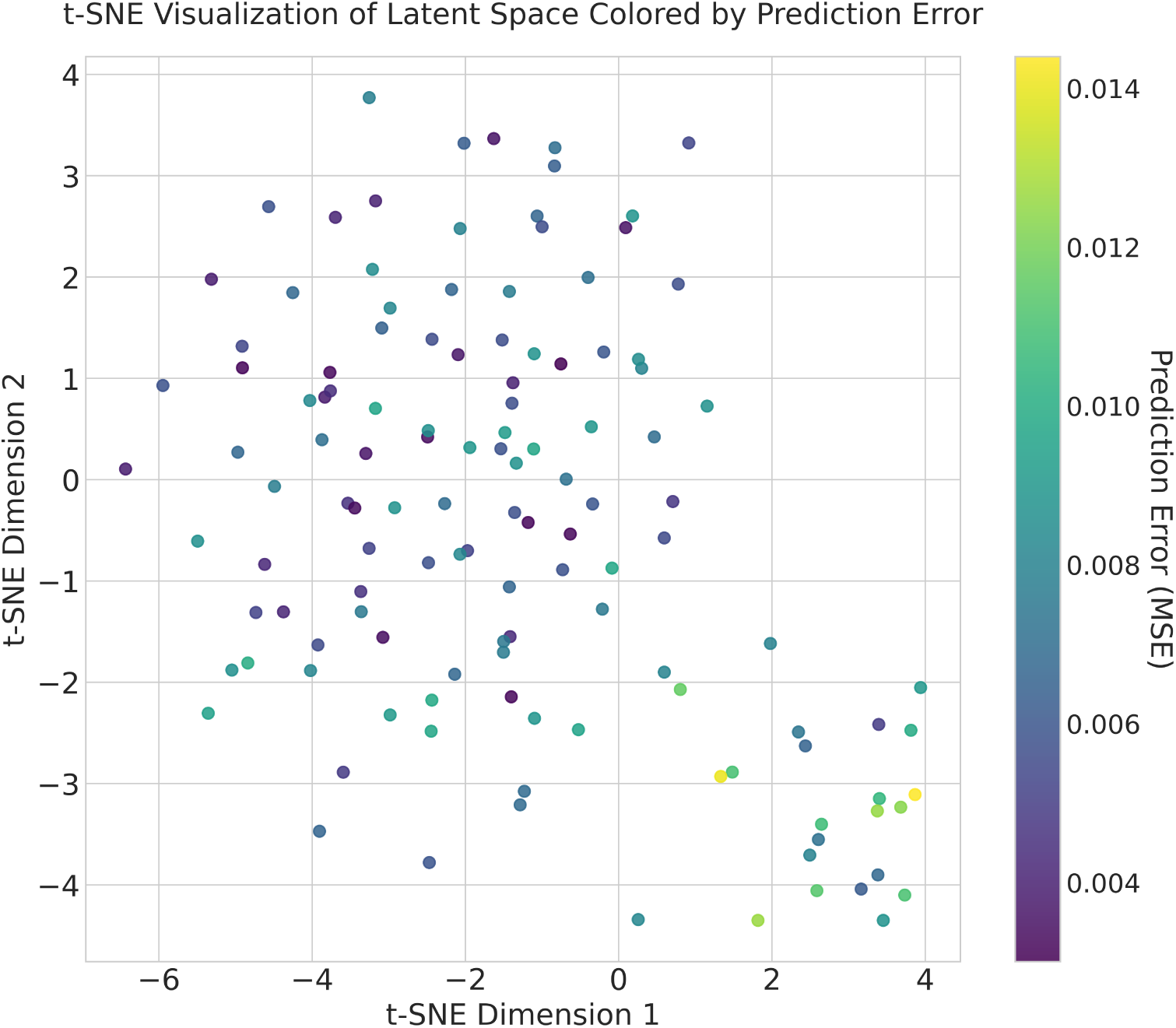
t-SNE analysis of the GMHA-AE model.

We observed some interesting relationships with age. The oFC-pFC correlation coefficients derived via the analytical model correlated negatively with healthy participant age for the non-NS weighted SC matrices (surviving FDR correction). The correlations were strongest for the alpha frequency band and the iRD-weighted SC matrices. This indicates that the analytical model is a better model of communication between brain areas for younger healthy participants than for older ones. It also points to the fact that alternative analytical models should be explored for describing communication between brain areas at different times across the life span. This result is in agreement with the results of an earlier study^22^, which reported decreases of the SC-FC relationship with age for fMRI-based FC. We also note that a similar trend was observed for a cohort of adolescent participants^23^. The fact that there is no correlation between participant age and oFC-pFC correlation coefficients when pFC is derived via the GMHA-AE model indicates that that model is more robust when it comes to detecting such relationships than the analytical model, and can capture intricate relationships even as those change over time.

We now discuss our results that pertain to psychosis. The GMHA-AE model that was trained on the healthy participant data, and which gave high prediction accuracy for those participants, did poorly at predicting the oFC of the psychosis participants, with the differences between the distributions of healthy and psychosis participants being highly statistically significant after FDR correction (Table 3). Despite the fact that the GMHA-AE model does not provide direct information about the communication mechanisms that underlie the SC-FC relationship, this finding clearly points to the fact that the features that the GMHA-AE model captures in the healthy-participant SC and FC matrices, and which result to it predicting oFC very well, are likely not present, or reduced, in the SC and/or FC connectomes of the psychosis participants.

In contrast to the poorer performance of the GMHA-AE model for the psychosis participants across frequency bands, the analytical model gave much better predictions for the FC of psychosis participants than that of healthy participants for the delta band, across SC edge weights. As the *p*-values for the differences between the healthy versus the psychosis participant distributions of the correlation coefficients indicate (Table 3), the difference is highly statistically significant. This indicates that the SPL-SI algorithm employed in our analytical model is a better representation of the communication mechanisms that are responsible for the evocation of delta band resting-state FC in psychosis than it is for healthy participants. When looking at the beta frequency band, however, the prediction accuracy of the same analytical model was lower for psychosis participants compared to healthy participants for the FA-, and FRt-weighted SC matrices; the same trend was present for the iRD-weighted SC matrices (*p* = 0.0307, but that *p*-value did not survive FDR correction). No statistically significant differences were observed in the correlation distributions between healthy participants and psychosis participants for the theta and alpha bands, with the exception of the {alpha band / FA} and {alpha band / iRD} combinations, where a reduction in the prediction accuracy was observed. These results are in agreement with the results reported in a different study^24^, where healthy participants exhibited larger correlation between the average functional connectivity (measured via fMRI) and white-matter microstructure in comparison to schizophrenia patients.

The performance of the two models on the psychosis participants depends strongly on the FC frequency band (Fig. 12). Specifically, the analytical model performs much better in the delta and theta frequency bands, while the GMHA-AE model performs better in the alpha and beta bands. Keeping in mind that the GMHA-AE model was trained on healthy participants, this implies that the healthy-participant communication mechanisms are worse at describing the SC-FC relationship in psychosis participants compared to the simpler analytical model based on the SPL-SI algorithms. For the alpha and beta frequency bands, on the other hand, the complex communication patterns that the GMHA-AE model captures from the healthy participants are better at describing the SC-FC relationship than the analytical model.

It is important to stress that, above and beyond the small FC-specific and SC-specific alterations that we observed in the psychosis participants ( Fig. 4 and 5), we observed alterations in the *relationship* between SC and FC which were much more pronounced, in particular when that relationship was predicted via the autoencoder model. Our study shows that investigating the SC-FC relationship goes further than the summary statistics presented in Fig. 4 and 5, and identifies differences in the *relationships* between the structural and functional connections.

Our study has some limitations. Firstly, the GMHA-AE model does not provide insight into the communication mechanisms between brain areas. This lack of insight is, however, compensated by the fact that the model can capture complex relationships that analytical algorithms of brain communication cannot fully incorporate. Future work will target the interpretability and explainability of the model. Secondly, even though the healthy participant sample we used was large (126 participants) the number of psychosis participants was low (5). This is an issue with many similar studies - in particular some psychosis participants who volunteered for our study were not eligible to be scanned in our state-of-the-art Connectom MRI scanner, resulting in a smaller sample than a study using more conventional scanners would have. However, the statistical tests we used when comparing healthy to psychosis participants are appropriate to use when there is such a difference in the two samples, and therefore the differences we detected between healthy and psychosis participants are robust. The high statistical significance of our results indicates the need for followup studies of larger cohorts.

To conclude, in this work, we proposed a brain SC-FC mapping framework based on the Graph Multi-Head Attention AutoEncoder. By employing an autoencoder, our GMHA-AE model is capable of efficiently handling the distributed and heterogeneous patterns of the brain and of learning complex relationships between brain structure and function by utilizing attention mechanisms. We compared the GMHA-AE performance to that of an analytical model. We also used both models to investigate the SC-FC relationship in psychosis participants and compare it to that of healthy participants, and identified crucial differences in that relationship between the two groups. In future, the use of deep-learning models could be explored to quantify the SC-FC relationship for younger age groups, for task-based electrophysiological recordings, as well as for neurological disorders other than psychosis.

## Supplementary Material

### 5 Data acquisition and Preprocessing

#### 5.1 Sample

Multi-modal (MRI and MEG) data were collected from healthy participants via the Welsh Advanced Neuroimaging Database (WAND) study^26^. In this work, we use data from those participants that have good quality MRI and MEG data. Under these conditions, our sample consists of 126 healthy participants between the ages of 18 and 50 (73 female), 101 of whom were between the ages of 18 and 35 (55 female). The age distributions of the participants are shown in Fig. 15). The same data were collected from 5 people with psychosis between the ages of 18 and 35 (3 female). All psychosis participants were diagnosed at least 9 months prior to data collection.

**Figure 15.**
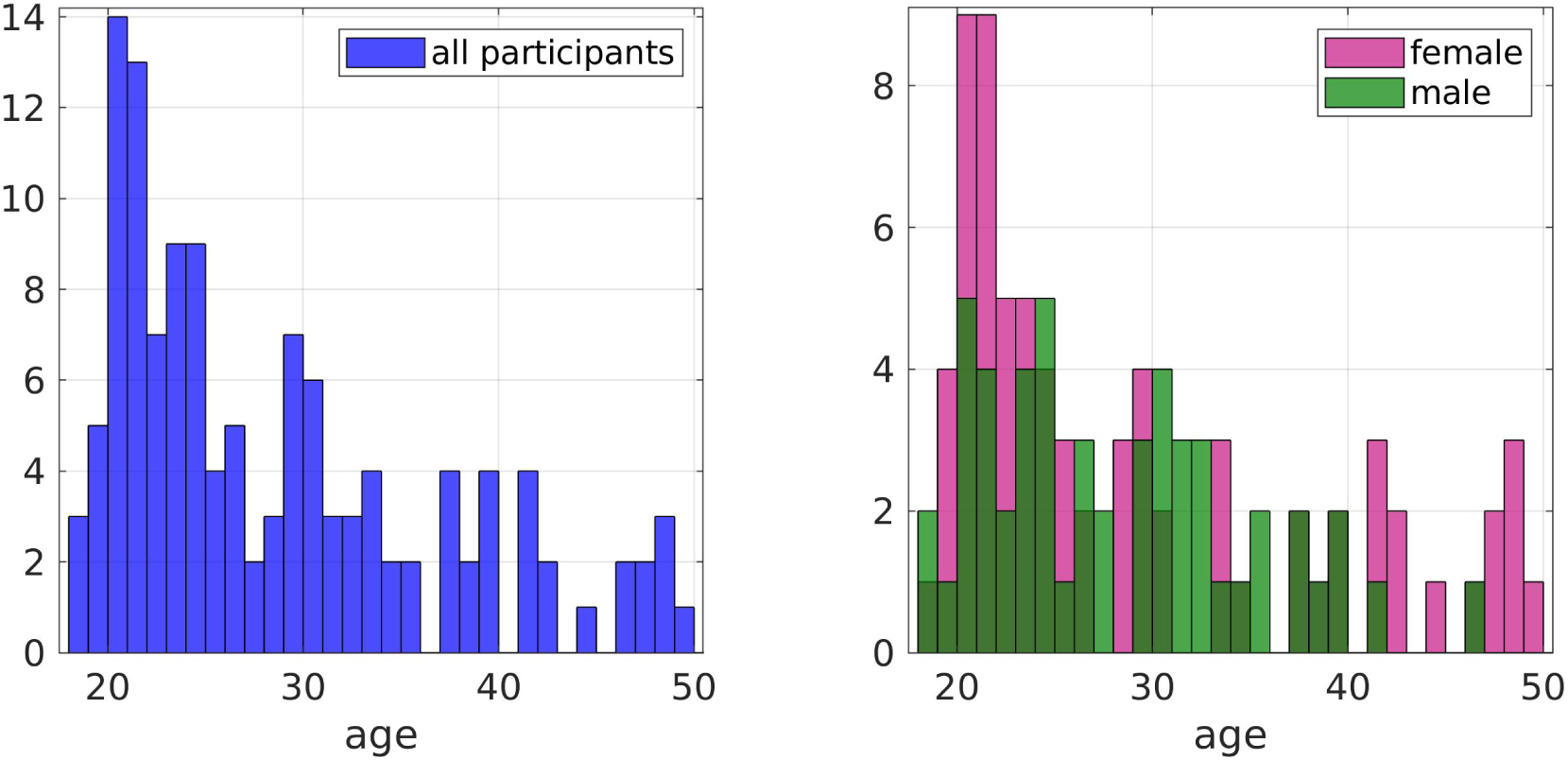
Age distributions of the healthy participants. The plot of the right panel breaks down the distribution by sex.

#### 5.2 MRI

##### 5.2.1 Data acquisition

All MRI data were acquired on a 3T Siemens Connectom scanner (300mT/m).

T1-weighted images were acquired using magnetization-prepared 180^◦^ radio-frequency pulses and rapid gradient-echo (MPRAGE) sequence, with repetition time (TR) 2300ms, echo time (TE) 2ms, field of view (FOV) 256 x 256 x 192 mm, matrix size 256 x 256 x 192, voxel size 1 x 1 x 1 mm, flip angle 9^◦^, inversion time (TI) 857ms, in-plane acceleration (GRAPPA) factor 2 and phase-encoding direction anterior to posterior (A»P).

Multi-shell diffusion-weighted MRI data were acquired using protocols described recently for the Microstructural Image Compilation with Repeated Acquisitions (MICRA) dataset^28^. Data were acquired over 18 minutes using a single-shot spin-echo, echo-planar imaging sequence, in both anterior to posterior (A»P) and posterior to anterior (P»A) phase-encoding directions. A»P data comprised two shells of 20 diffusion encoding directions uniformly distributed (Jones et al., 1999) at b = 200 s/mm^2^ and b = 500 s/mm^2^, one shell of 30 directions at b = 1200 s/mm^2^ and three shells of 61 directions each at b = 2400 s/mm^2^, 4000s/mm^2^ and 6000 s/mm^2^. Additionally, two leading non-diffusion-weighted (b = 0 s/mm^2^) images and 11 non-diffusion-weighted images were acquired, dispersed throughout (33^rd^ volume and every 20^nd^ volume thereafter). P»A data comprised two leading non-diffusion-weighted images, one shell of 30 directions at b = 1200 s/mm^2^ and a final non-diffusion-weighted image. Data acquisition details for all b-values are as follows: TR=3000ms, TE=59ms, FOV = 220 x 200 mm in-plane, matrix size 110 x 110 x 66, voxel size 2 x 2 x 2 mm, with in-plane acceleration (GRAPPA) factor 2. The diffusion gradient duration and separation were 7ms and 24ms, respectively.

##### 5.2.2 Data analysis

Data were analyzed on a CentOS Linux 7 cluster computing system.

Cortical reconstruction and volumetric segmentation of the T1-weighted images was performed with the FreeSurfer image analysis suite (https://surfer.nmr.mgh.harvard.edu). The technical details of the methods used by the FreeSurfer software have been described in other publications^29–41^. The Desikan-Killiany atlas^61^ was used to identify the grey matter areas that form the nodes of the structural brain networks. The atlas provides 82 cortical and subcortical areas for the left and right cerebrum.

The diffusion-MRI data were corrected for thermal noise, signal drift, susceptibility distortions, motion and eddy-current distortions, gradient non-uniformity and Gibbs ringing artifacts using a combination of in-house pipelines, the FMRIB Software Library^54^ and MRtrix3^45^. A brain mask was created using the Brain Extraction Tool^54^ to exclude non-brain data. Diffusion MRI noise level estimation and denoising was performed using a Marchenko-Pastur principal component analysis (MP-PCA)- based approach^83,84^. Within-image intensity drift was corrected by fitting the diffusion-MRI data to temporally interspersed b0 images using in-house code in MATLAB R2017b (MathWorks Inc. Natick, Massachusetts, USA). Slicewise OutLIer Detection (SOLID)^85^ was applied, with lower and upper thresholds of 3.5 and 10 respectively, using a modified z-score and a variance-based intensity metric. The susceptibility-induced off-resonance field was estimated from the b0 data collected in opposing phase-encoding directions using FSL’s topup^42,43^ and corrected, along with eddy-current induced distortions and participant movement using FSL’s eddy tool^44^. In-house code was used to correct for gradient non-uniformity distortions in MATLAB 2017b (MathWorks Inc. Natick, Massachusetts, USA). Gibbs ringing correction was performed in MRtrix3^45^ using the subvoxel-shifts method^58^.

MRtrix3^45^ was used to calculate the response function using the Dhollander algorithm^46,47^, and the fiber orientation distributions using Multi-Shell-Multi-Tissue constrained spherical deconvolution^52^, in order to perform anatomically-constrained tractography. The MRtrix function 5ttgen^43,48–51^ was used to segment anatomical images into cortical grey matter (GM), subcortical grey matter, cerebrospinal fluid, white matter (WM) and abnormalities. Anatomical images were coregistered to the diffusion-weighted images using FSL^43,53–56^. The interface between grey and white matter was identified using function 5tt2gmwmi of MRtrix3^45,49^. Anatomically-constrained streamline tractography^45,49,86^ was used to generate whole-brain tractograms with seeds in the GM-WM interface. Streamlines were of minimum/maximum length of 30/250mm, cutoff of 0.06, and maximum angle between successive steps of 50^◦^. Twenty million streamlines were generated for each participant. The sift2 algorithm^57^ was used to provide tractograms that have a density of reconstructed connections proportional to the fibre density within each voxel as estimated by the diffusion model. This ensures that the number of streamlines connecting two regions of grey matter provides an estimate of the cross-sectional area of the white matter axons connecting those regions, which is a biologically-relevant measure of structural connectivity. Each participant’s tractogram was overlayed on the participant’s fractional anisotropy map and visually inspected to ensure that the tractogram provided good coverage of the white matter and that no streamlines extended into unphysical regions.

The structural connectivity matrices representing the structural networks of the participants have the cortical and subcortical areas identified from the Desikan-Killiany atlas as the nodes, and the white matter tracts linking those areas as the edges (connections). To evaluate the impact that different white matter characteristics have on predicting electrophysiological resting-state FC, we used the following 5 structural measures as edge weights in the SC matrices.

1. Number of reconstructed streamlines (NS) connecting two cortical regions: SC matrices weighted with NS are good at predicting functional connectivity from both fMRI^2,10^ and MEG^17^.
2. Volume-normalized NS (NS/v): Because the absolute number of streamlines of a connection depends on the size of the connected brain areas (i.e., pairs of larger brain areas are connected by more streamlines because of their size rather than because of stronger connectivity) we divided the NS by the sum of the volumes of the connected brain areas.
3. Fractional anisotropy (FA) of the diffusion tensor: The FA is related to the myelination and axonal characteristics of white matter tracts, and therefore is a good proxy for structural connectivity^10,17,64–66^.
4. Inverse radial diffusivity (iRD) of the diffusion tensor: The iRD is related to the myelination of the white matter tracts, and has been used in studies in health and disease^17,70^.
5. Total restricted signal fraction (FRt): derived from the CHARMED model^71^, attributed to water within the intra-axonal space and therefore related to axonal characteristics of the WM tracts.

Our choice to investigate the FRt is motivated by the non-specificity of the FA and iRD to biological attributes of the brain’s WM, such as myelination and axonal diameter. For example, a reduction in FA could reflect reduced neurite density, increased neurite orientation dispersion, both, or various other changes to tissue microstructure^87^. Radial diffusivity can be influenced by both myelination and axonal density of the white matter tracts, and therefore higher values can indicate lower myelination or axonal density^67–69^.

With these 5 measures as edge weights (NS, NS/v, FA, iRD, FRt) we have 5 SC matrices for each participant. These 5 matrices all represent the participant’s structural substrate that underlies FC, but each relies on a different aspect of the brain’s microstructure to depict structural connectivity strength. The SC matrices were normalized so that the values for each SC matrix of each participant were in the range of 0 to 1.

The Euclidean distance (ED) between centers of the Desikan-Killiany brain areas was calculated for all participants. Specifically, MRtrix^45^ was used to locate the coordinates of the center of each brain area, and in-house Matlab code was used to calculate the distances between those. Structural matrices were then calculated in which the values of structural connections were the Euclidean distances between the centres of the DK brain areas.

#### 5.3 MEG

##### 5.3.1 Data acquisition

Ten-minute whole-head MEG recordings were acquired in a 275-channel CTF radial gradiometer system, at a sampling rate of 1,200 Hz. Twenty-nine additional reference channels were recorded for noise cancellation purposes and the primary sensors were analyzed as synthetic third-order gradiometers^88^. Participants were seated upright in a magnetically shielded room with their head supported with a chin rest to minimize movement. They were asked to rest with their eyes open and to fixate on a central red point presented on a PROPixx LCD projector (Vpixx Technologies Inc). Horizontal and vertical electro-oculograms (EOG) were recorded to monitor eye blinks and eye movements. Recordings were also acquired while the participants performed tasks after the completion of the resting-state paradigm, but those recordings were not used in the analysis presented here.

###### Data analysis

To achieve MRI/MEG co-registration, fiduciary markers were placed at fixed distances from three anatomical landmarks identifiable in the participant’s T1-weighted anatomical MRI scan, and their locations were manually marked in the MR image. Head localization was performed at the start and end of the MEG recording and continuously through the recording. The data were subsequently pre-processed in a manner similar to that described in previous work^60^. Specifically, all datasets were downsampled to 600 Hz, and filtered with a 1 Hz high-pass and a 150 Hz low-pass filter. The datasets were then segmented into 2-s epochs, and were visually inspected. Epochs exhibiting large head movements, muscle contractions, or other large artefacts were excluded from subsequent analysis.

The functional networks were constructed in a manner similar to that described in earlier work^60^ but with some important differences. Specifically, the MEG sensor data were downsampled to 300 Hz, before being source localized using FieldTrip (RRID:SCR_004849) version 20190219^59^. The source-reconstruction grid for each participant was constructed using Freesurfer to segment the individual’s T1 anatomical scan to yield the set of gray-matter voxels at a resolution of 2 mm isotropic. All these voxels were automatically labelled using the Desikan-Killiany parcellation^61^ and matched to the MEG-coregistered MRI for the individual using FSL-FLIRT. All of these coregistered and parcellated gray-matter voxels were used for LCMV beamformer source-localisation using a single-shell forward model^89^, where the covariance matrix was constructed in each of the four lower frequency bands: delta (1–4 Hz), theta (4–8 Hz), alpha (8–13 Hz), and beta (13–30 Hz). For each band, the beamformer weights were normalized using a vector norm^90^. Epochs were concatenated to generate a continuous dataset that was projected through these weights to yield a virtual-sensor time course for each voxel and then band passed to the above-mentioned frequency bands. For each of the 82 Desikan-Killiany regions, the virtual channel at the centroid of the region was chosen as a representative voxel.

The resulting 82 time series were orthogonalized using symmetric orthogonalization^91^, to avoid spurious correlations. A Hilbert transform was used to obtain the oscillatory amplitude envelope. The data were subsequently despiked using a median filter in order to remove artifactual temporal transients, downsampled to 1 Hz, and trimmed to avoid edge effects (removing the first two and the last three samples). To derive the amplitude-amplitude connectivity, amplitude correlations were calculated by correlating the downsampled Hilbert envelopes to each other, and the resultant correlation coefficients were converted to variance-normalized z-scores. This choice was motivated by the fact that such correlations have been shown to be one of the most robust and repeatable electrophysiological connectivity measures^92,93^. Specifically, these studies found that the oscillatory amplitude envelope correlation gave the most consistent connectivity, in contrast to other methods which lack repeatability over scans of the same subject. Given that we are comparing healthy participants to a patient population, it is important to have the most robust measure of functional connectivity possible.

Through this process we obtained 4 FC matrices for each participant.

### 6 Autoencoder Details

Based on the effectiveness of graph networks for representing brain topological structures, the encoder and decoder in this study are both selected to use graph network structures.

#### 6.1 Encoder Structure

The graph attention mechanism is employed as the encoder layer to generate effective high-dimensional latent feature representations. By adding multiple layers of graph attention mechanisms to the encoder structure, the model’s depth and learning capability are enhanced. Additionally, the representation learning of graph structure nodes produces richer node embeddings. To determine the relation between nodes and their neighbors, we use a self-attention mechanism that shares parameters between nodes. The input required by the encoder is the symmetrical adjacency matrix of the structural connectivity *A* ∈ R*^n^*^×*n*^ and the set of node features *x* = {*x*_1_*, x*_2_*, . . . , x_n_*}, where *n* = 82 represents the number of nodes, i.e., the number of cortical regions partitioned in the DK brain atlas. The initial feature of node *i* is *h*^0^ = *x_i_* ∈ R^1×*d*^. In the *l*-th layer of the encoder, the attention coefficient between node *i* and its neighbor node *j* is calculated as follows:

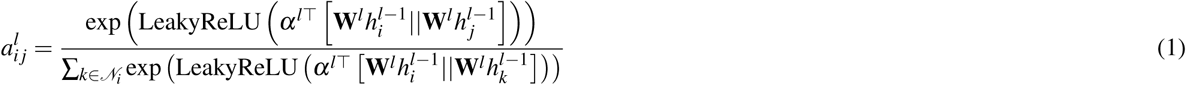

where (*N_i_*) is a set of nodes connected to node *i* according to the adjacency matrix *A*, *α* denotes a learnable weight vector, || represents the concatenation operation, **W** is a learnable linear transformation weight matrix, and LeakyReLU is the activation function. Further, the output representation of node *i* in the *l*-th layer of the encoder is as follows:

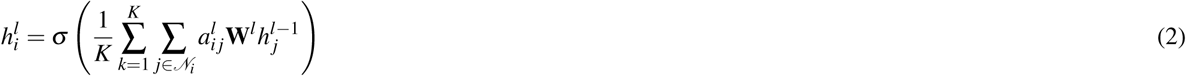

After passing through *L* encoder layers, the output of the last layer is regarded as the final node embedding, i.e., *h_i_* = *h_i_^L^,* ∀*i* ∈ {1, 2*, . . . , n*}.

#### 6.2 Decoder Structure

Contrary to the "compression" in the encoder, the decoder’s role is to reconstruct the high-dimensional representations into the size of the functional connectivity matrix. We use a decoder with the same number of layers as the encoder, where each decoder layer reverses the process of its corresponding encoder layer, i.e., each decoder layer reconstructs the representation of the nodes by utilizing the representations of their neighboring nodes (according to their relevance). In the *l*-th decoder layer, the attention coefficient between node *i* and its neighbor node *j* is calculated as follows:

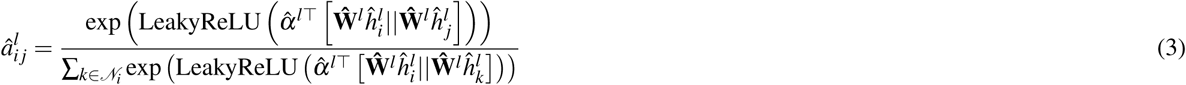

Similar to Equation 1, **Ŵ** and *α̂* are learnable weight matrices.

Taking the output of the encoder as the input to the decoder, i.e., 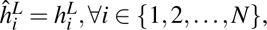 the representation of node *i* in the (*l* − 1)-th layer of the decoder is reconstructed as follows:

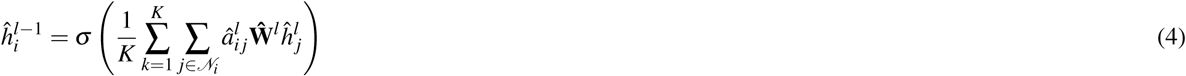

After passing through the same *L* layers of the decoder, the set of outputs of all nodes in the last layer is regarded as the reconstructed functional connectivity matrix, i.e., 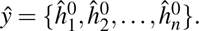

#### 6.3 Objective Function

To achieve accurate predictions of functional connectivity from the structural connectivity matrices, the objective function of the neural network is designed to meet two requirements: 1) minimizing the prediction error between the oFC and the model-predicted pFC, which is achieved by calculating the mean squared error between oFC and pFC, 2) introducing a regularization term to ensure that the correlation (inter-pFC) between different individuals in the pFC is comparable to the correlation (inter-oFC) between different individuals in the oFC. This ensures that the neural network learns the mapping from SC to oFC while retaining individual differences, rather than predicting a population-average representation of functional connectivity for all individuals. Based on these two requirements, the objective function for the proposed overall structure is:

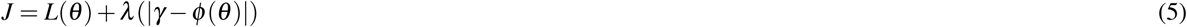

where *θ* represents the parameters of the neural network to be learned, *L*(·) is the loss function, and *λ* is a regularization constant used in the regularization function |*γ* − *ϕ* (·)|, similar to the L1 norm. The mean squared error is used as the loss function, as shown below:

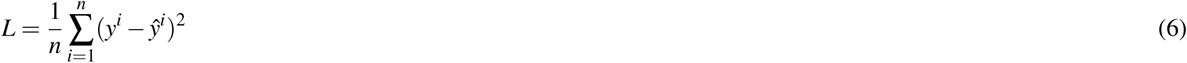

where *n* is the number of samples in a training batch, *i* represents a sample, *y* is the actual label (oFC), and *y*^ is the predicted output (pFC). The mean squared error ensures that the spatial correlation between SC and FC is approximately preserved during training.

The objective of the overall autoencoder structure is to predict the actual FC while also maintaining the inter-oFC differences between individuals. Therefore, a regularization term is added to the objective function, and the Pearson correlation coefficient *r* is introduced into the regularization function |*γ* − *ϕ* (*θ*)|:

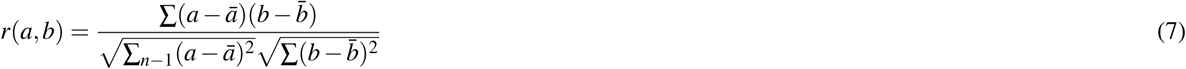

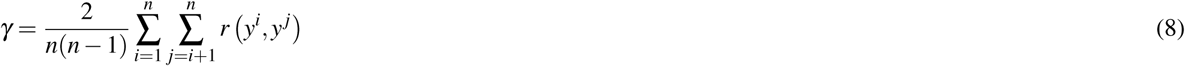

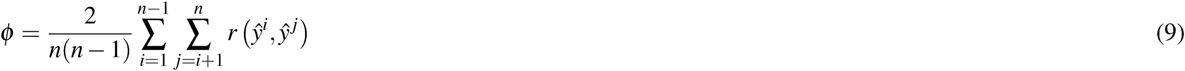

In the above equations, *γ* represents the inter-oFC correlation of the training dataset, which remains unchanged during training. It is noteworthy that *γ* varies for different datasets. In this case, *ϕ* represents the inter-pFC correlation, which is expected to be less than or equal to the inter-oFC correlation. Thus, the regularization function *λ* (|*γ* − *ϕ* (*θ*)| ensures that the inter-individual differences in the pFC are comparable to the inter-individual differences in the oFC.

### 7 Performance Variability

To further assess the robustness of the model’s outputs across subjects, we evaluated the stability of the correlation-based features over 5-fold cross-validation. Each fold contained 26, 25, 25, 25, and 25 subjects, respectively. For each feature matrix (NS, NS/v, FA, iRD, FRt), we computed the average value across subjects within each fold, and then measured the cross-fold standard deviation and 95% confidence intervals. The results are summarized in Table below. Notably: the alpha and beta bands consistently show high stability, with standard deviations <0.02 and CI widths <0.04 across all feature types. The delta and theta bands show higher variability.

## 8 Data availability

The WAND data is freely available at: https://doi.org/10.12751/g-node.5mv3bf.

## 10 Acknowledgements

We acknowledge contributions of Svetla Manolova, Elena Stylianopoulou, Emma Morgan, Gavin Perry, Alexander Shaw, Mark Drakesmith and Holly Rossiter to the WAND data collection. We also wish to thank all the participants of the WAND study.

**Table 5.** Performance variability measures.

| SC metric | FC frequency band | Mean | SD | 95% CI |
| --- | --- | --- | --- | --- |
| NS | delta | 0.13 | 0.12 | [-0.01, 0.28] |
|  | theta | 0.50 | 0.02 | [0.48, 0.53] |
|  | alpha | 0.83 | 0.01 | [0.81, 0.85] |
|  | beta | 0.85 | 0.01 | [0.84, 0.87] |
| NS/v | delta | 0.09 | 0.16 | [-0.11, 0.29] |
|  | theta | 0.48 | 0.03 | [0.45, 0.52] |
|  | alpha | 0.84 | 0.02 | [0.81, 0.86] |
|  | beta | 0.85 | 0.01 | [0.84, 0.87] |
| FA | delta | 0.34 | 0.17 | [0.12, 0.55] |
|  | theta | 0.49 | 0.18 | [0.27, 0.72] |
|  | alpha | 0.86 | 0.01 | [0.84, 0.87] |
|  | beta | 0.86 | 0.01 | [0.85, 0.87] |
| iRD | delta | 0.42 | 0.04 | [0.37, 0.48] |
|  | theta | 0.51 | 0.06 | [0.44, 0.58] |
|  | alpha | 0.86 | 0.01 | [0.85, 0.87] |
|  | beta | 0.86 | 0.01 | [0.85, 0.87] |
| FRt | delta | 0.50 | 0.01 | [0.48, 0.52] |
|  | theta | 0.48 | 0.14 | [0.31, 0.66] |
|  | alpha | 0.84 | 0.03 | [0.81, 0.87] |
|  | beta | 0.86 | 0.01 | [0.84, 0.87] |

## 11 Author contributions statement

QC: Setup of the autoencoder, data analysis, writing of the manuscript.

HT, VH: Data collection, data curation, data analysis.

PLL: Analysis pipeline development.

CBM: Data collection, data analysis.

KDS: Data analysis, writing of the manuscript.

DKJ: Study design, writing of the manuscript.

EM: Data collection, data analysis, study design, writing of the manuscript. All authors reviewed the manuscript.

